# Fast and accurate bacterial species identification in biological samples using LC-MS/MS mass spectrometry and machine learning

**DOI:** 10.1101/635227

**Authors:** Florence Roux-Dalvai, Clarisse Gotti, Mickaël Leclercq, Marie-Claude Hélie, Maurice Boissinot, Tabiwang N. Arrey, Claire Dauly, Frédéric Fournier, Isabelle Kelly, Judith Marcoux, Julie Bestman-Smith, Michel G. Bergeron, Arnaud Droit

## Abstract

The identification of microbial species in biological samples is essential to many applications in health, food safety and environment. MALDI-TOF MS technology has become a tool of choice for microbial identification but it has several drawbacks including: it requires a long step of bacterial culture prior to analysis (24h), it has a low specificity and is not quantitative. We have developed a new strategy for identifying bacterial species in biological samples using specific LC-MS/MS peptidic signatures. In the first training step, deep proteome coverage of bacteria of interest is obtained in Data Independent Acquisition (DIA) mode, followed by the use of machine learning to define the peptides the most susceptible to distinguish each bacterial species from the others. Then, in the second step, this peptidic signature is monitored in biological samples using targeted proteomics. This method, which allows the bacterial identification from clinical specimens in less than 4h, has been applied to fifteen species representing 84% of all Urinary Tract Infections (UTI). More than 31000 peptides in 200 samples have been quantified by DIA and analyzed by machine learning to determine an 82 peptides signature and build prediction models able to classify the fifteen bacterial species. This peptidic signature was validated for its use in routine conditions using Parallel Reaction Monitoring on a capillary flow chromatography coupled to a Thermo Scientific™ Q Exactive HF-X instrument. Linearity and reproducibility of the method were demonstrated as well as its accuracy on donor specimens. Within 4h and without bacterial culture, our method was able to predict the predominant bacteria infecting a sample in 97% of cases and 100% above the 1×10^5^ CFU/mL threshold commonly used by clinical laboratories. This work demonstrates the efficiency of our method for the rapid and specific identification of the bacterial species causing UTI and could be extended in the future to other biological specimens and to bacteria having specific virulence or resistance factors.

## INTRODUCTION

The identification of the bacterial species or strain present in a biological sample is essential in many fields of microbiology. Epidemiology, for instance, tracks the spreading of microorganisms related to infectious diseases; food safety laboratories ensure the distribution of pathogens-free products to the consumers; environmental bacteria have a strong impact on maintaining the equilibrium of ecosystems; and clinical laboratories require fast diagnosis methods to provide appropriate treatment to patients with a bacterial infection. However, standard methods for the identification of pathogens requires a time-consuming bacterial culture followed by another long step of immunological or biochemical tests of varying duration and cumbersomeness (1–3). During this period, typically of 24 to 48h but could extend to weeks, patients received broad spectrum antimicrobial treatments which might be not optimal and sometimes not even efficient to fight the infection (4). Furthermore, inappropriate use of antimicrobial agents contributes to the selection the resistant bacteria in the whole population thus making infections more challenging to combat (5, 6). Therefore, there is a need for the development of fast and robust methods for bacterial identification, in order to improve therapy and guide rational use of antibiotics.

Genotyping methods, which are based on the sequencing of partial (16S small subunit ribosomal [rRNA] gene sequencing) or entire genomes (Whole Genome Sequencing) of the microorganisms contained in a sample, are promising since they do not require bacterial culture and can be applied to complex samples containing several species (7, 8). However, the cost and the time required to get identification by sequencing methods preclude their use in routine laboratories. In addition, if 16S rRNA sequencing can provide a quite rapid identification (typically 24 hours), the high conservation of 16S gene sequences across bacterial families and species often limits the precision of identification to the genus level (9, 10). By contrast, Whole Genome Sequencing is able to provide an efficient species and even strain typing, but the cost and the time required to get the results is strongly extended by the sequencing itself and by the data analysis. Moreover, this analysis requires expert scientific knowledge to provide a confident genome assembly as well as large computing resources (11, 12).

In the past few years, Matrix-Assisted Laser Desorption Ionization – Time Of Flight Mass Spectrometry (MALDI-TOF MS) analysis of microbial proteins has made a breakthrough in routine labs for bacterial identification (13–16). This fast, inexpensive, and automatable technology can replace the conventional phenotype-based methods, hence reducing the time required to get an identification from 2 or 4 days to less than 50 hours. For those reasons, two mass spectrometers, the Biotyper (Bruker) and the Vitek-MS (Shimadzu-BioMérieux), have been approved for clinical use by health governmental organizations of most countries including the United States Food and Drug Administration (FDA) in 2013 (17). In the typical workflow, bacterial colonies isolated by culture are submitted to fast sample preparation (typically, a treatment with formic acid and ethanol) prior to acquisition of protein mass spectra that are used to interrogate a spectral database providing a confidence score for the bacterium identification, an information a physician can use to diagnose the infection.

Despite its numerous advantages, bacterial identification by MALDI-TOF MS has several drawbacks: i) it requires a lengthy culture step to isolate bacterial colonies, since the detection is based on a comparison with spectral database acquired on pure colonies. For the same reason, is not able to identify polymicrobial infections (*i.e.*: when several species are present in the same sample) without analyzing several types of colonies visually selected on the culture plate; ii) because of the minimal sample preparation, the information contained in the spectra is restricted to the most abundant molecules, thus limiting the specificity of the method and its capability to identify certain species or subspecies and; iii) it is not quantitative, a potentially important information for certain specimens where pathogens need to be distinguished from the normal microbiota, or when a certain level of infection needs to be reached to necessitate antibiotherapy.

To overcome the abovementioned issues, several studies have tried to improve MALDI-TOF bacterial identification (18). For instance, Clark and colleagues refined the specificity of the method to identify *Escherichia coli* pathotypes by examining specific peaks in the spectra (19). Other investigators have tried to improve the specificity using trypsin digestion which allows the accession to a larger set of molecules and the generation of a Peptide Mass Fingerprint of the bacterial subspecies (20). Several studies skip the culture step to provide a faster identification, especially in the case of sepsis where MALDI-TOF acquisition is performed directly from a positive blood culture sample (21, 22). However, it has been shown that sample preparation methods, which are not homogenous from lab to lab, can influence the rate of correct identifications of certain microorganisms (23). Although these studies could improve the standard workflow, they are limited by the sensitivity and the specificity of MALDI-TOF mass spectrometers. Therefore, recent studies have investigated the possibility of using LC-MS (Liquid Chromatography-Mass Spectrometry) methods which, because of their high sensitivity and specificity, have replaced MALDI-TOF MS in most research laboratories. Wang and colleagues used the LC-MS approach to identify biomarkers of five major bacterial species in brochoalveolar lavage specimen (24) and performed strain typing for *Acinobacter baumanii* (25), Karlsson R et *al.* used it for proteotyping within the mitis group of *Streptococcus* genus (26) and Cheng *et al.* also used LC-MS/MS in Selected Reaction Monitoring (SRM) mode to target specific peptides of the flagella to type *Escherichia coli* at strain level (27). Bioinformatical tools have also been developed to help in the identification of bacteria from ‘bottom up’ proteomics data (*i.e.* trypsin-digested proteins). These methods were able to reach 89 to 98.5% correct classification rates at the species level but these values have only been demonstrated after a step of bacterial growth (28, 29).

Taking the advantages of sensitivity and specificity from nanoscale LC-MS/MS technology, and based on these previous studies, we developed a new pipeline using modern proteomics (DIA – Data Independent Acquisition mode) and machine learning algorithms to identify biomarkers able to speciate a set of bacteria of interest. This strategy is based on two steps (Figure 1): i) a training step, that enables to define a peptidic signature for the bacteria of interest and ii) an identification step where the signature is monitored by targeted proteomics to get the identification of the bacteria in the infected samples. Once the training step has been developed, the second step can be performed in routine laboratories on multiple samples and with any type of mass spectrometer working in PRM (Parallel Reaction Monitoring) or SRM (Selected Reaction Monitoring) modes.

**Figure 1.**
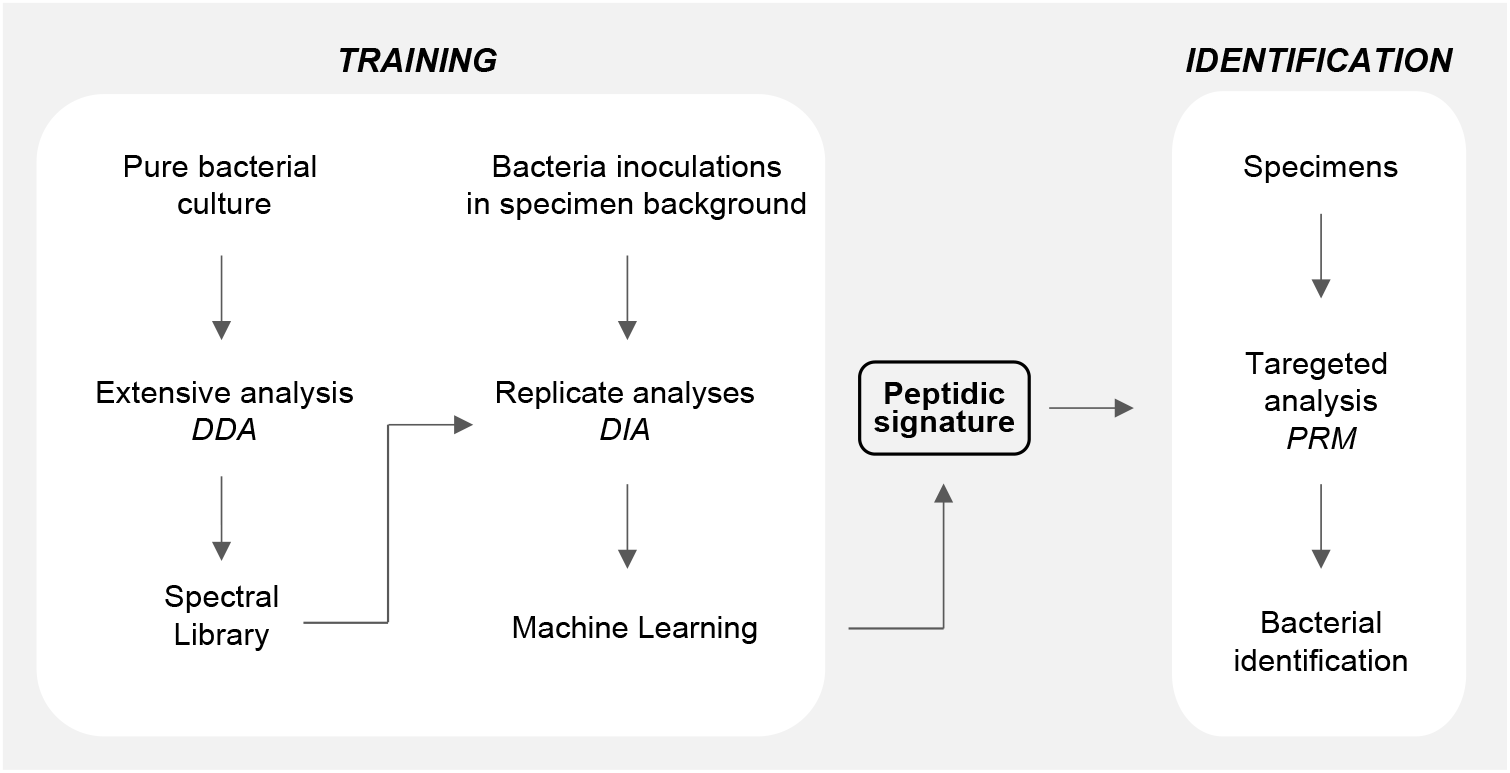
Workflow of the method for bacterial identification. The workflow is composed of two steps: the “training” step defines of a peptidic signature for the bacteria of interest; the “identification” step uses this signature in routine to identify bacteria in biological samples.

As a proof of concept, this pipeline has been applied to the 15 bacterial species most frequently found in Urinary Tract Infections (UTI). Indeed, urine is the most common clinical specimen with hundreds of samples analyzed each day in most of clinical laboratories. Moreover, UTI is one of the most frequent type of infection in humans: it has been demonstrated that 50 to 60 % of women in western countries will have at least one UTI in their lifetime (30). As reported by statistics of the Enfant-Jésus hospital in Québec City, which analyzes 300 urine specimens each day on average, 68.2% of these samples are infected by the same 4 bacterial species (*Escherichia coli, Streptococcus agalactiae, Klebsiella pneumoniae* and *Enterococcus faecalis*) and 15 species are responsible for more than 84% of all UTI (Supplementary Figure 1). According to literature reports, these are the most frequently found species in UTI (30, 31).

Our original method enables to define a peptidic signature which, when monitored by targeted proteomics, is able to detect which of the 15 bacterial species is present in the urine sample, in less than 4 hours, without any bacterial culture. We also demonstrated that the peptidic signature is transferable to other laboratories and to other mass spectrometers. In addition, we compared the efficiency of our method to the MALDI-TOF standard workflow.

## EXPERIMENTAL PROCEDURES

### Bacterial culture and counting

Bacterial strains were obtained from the Culture Collection of Centre de Recherche en Infectiologie of Université Laval (CCRI, Québec, Canada).

The bacterial strains used and their corresponding culture conditions are listed in supplementary methods. Semi-log broth bacterial culture calibrated to 0,5 MacFarland suspension were prepared and used for spectral libraries generation or for urine inoculation. In parallel, they were counted by incubation of 100uL of serial dilutions on blood agar plate (Supplementary Methods).

### Urine collection and bacterial inoculation

To mimic urinary tract infections, 1 to 200 µL of semi-log broth culture suspension, corresponding to an estimated final amount of 1×10^4^ to 1×10^6^ CFU/mL, were spiked into 10mL of urine obtained from six different healthy volunteers. The exact concentration of the inoculated cultures was determined in parallel by culture on agar plates as described above.

Moreover, urine specimens from 27 patients were collected at the Microbiology-Infectiology laboratory of Enfant-Jésus hospital of CHU de Québec (Québec, Canada) after few microliters had been used for standard MALDI-TOF analysis. The specimens were kept on ice during the transportation (< 1h) and used immediately.

The consent of all donors was obtained as described in the ethical approval of the Comité d’Éthique de la Recherche of CHU de Québec – Université Laval (recording number 2016-2656).

### Sample preparation for spectral libraries

For the generation of bacterial spectral libraries, bacteria from 1mL of semi-log broth bacterial culture were pelleted by centrifugation at 10,000 × *g* for 15 minutes, the supernatant was discarded and the pellet was washed three times with 1mL of 50mM Tris and centrifuged in the same conditions. The final pellet was frozen dried and stored at −20°C.

Pellets were then resuspended with 50mM of ammonium bicarbonate and 600 units of mutanolysine (Sigma-Aldrich, cat no. M9901) were added to help bacterial lysis by digestion of cell wall peptidoglycan. After a 1-hour incubation at 37°C, 0.5% sodium deoxycholate (SDC) and 20mM dithiothreitol (DTT) (final concentrations) were added and bacterial inactivation was performed by heating 10min at 95°C. Lysis was achieved by sonication for 15min with a Bioruptor® system (Diagenode), with cycles of 30s ON/30s OFF, high level. A final centrifugation at 16,000 × *g* during 15min was performed to remove cell debris, and protein concentration in the supernatant was measured using a Bradford assay.

Prior to proteolytic digestion, SDC concentration was adjusted to 1% and 120 µg of proteins from each bacterial culture were digested by addition of trypsin (Promega) in a 1:50 (enzyme:protein) ratio, during 1 hour at 58°C. Trypsin reaction was then stopped by acidification with 350µL of 5% formic acid (FA), which also leads to precipitation of the SDC. After centrifugation at 16,000 × *g* for 5 min, the supernatant was collected, the peptides were purified on Oasis HLB cartridge 10mg (Waters) and vacuum-dried.

The pellet was resuspended in 10mM Ammonium bicarbonate pH10 and an equivalent of 110µg of peptides were fractionated on an Agilent 1200 Series System HPLC equipped with Agilent extend C_18_ (1.0mm × 150mm, 3.5µm) column. Peptides were loaded at 1mL/min of solvent A (10mM ammonium bicarbonate pH10) and eluted by addition of solvent B (90% acetonitrile, 10% ammonium bicarbonate pH10) with a gradient 5 to 35% solvent B during 60 min and 35 to 70% solvent B during 24 min. Fractions were collected in a 96 well plates at 1 min intervals and finally pooled in rows into 8 fractions which were vacuum-dried.

Each fraction was resuspended in 2% acetonitrile (ACN) / 0.05% trifluoroacetic acid (TFA) at 0.2 µg/µL and 1X iRT peptides (Biognosys) were added. An equivalent of 1µg of peptides was injected on LC-MS/MS system for each fraction of each bacterial species.

### Preparation of urine samples

Urine specimens (10 mL), either from patients or artificially inoculated from healthy urine, were treated the same way: human cells were initially pelleted by a low speed centrifugation for 5 min at 1,000 × *g*, and the supernatant was high speed centrifuged for 15 min at 10,000 × *g* in order to collect bacteria. Bacterial pellets were then washed with 1mL of 50mM Tris and centrifuged again in the same conditions, another cycle of wash and centrifugation was added and the resulting pellet was frozen dried.

Protocols for protein extraction, trypsin digestion and peptide purification are described above in the ‘Sample preparation for spectral libraries’ section and were modified as follows: for each sample, 50 units of mutanolysine was used, 250 ng of trypsin was added for the digestion which was then stopped with 1µL of 100% FA and peptides were purified with StageTips (32) containing C18 reverse phase (3M Empore C18 Extraction Disks). Samples were resuspended in 10μL of 2% ACN, 0.05% TFA and 1X iRT peptides (Biognosys) were added. Half of the final volume was injected on LC-MS/MS system.

### LC-MS/MS acquisitions

Samples were analyzed by nanoLC/MS using a UltiMate^TM^ 3000 NanoRSLC system (ThermoScientific, Dionex Softron GmbH, Germering, Germany) coupled to an Orbitrap Fusion Tribrid – ETD mass spectrometer (ThermoScientific, San Jose, CA, USA, Instrument Control Software version 2.0). Peptides were trapped at 20 μL/min in loading solvent (2% acetonitrile, 0.05% TFA) on a μ-Precolumn, 300 μm i.d × 5mm, C18 PepMap100, 5 μm, 100Å (Thermo Fisher Scientific) during 5 minutes. Then, the pre-column was switched online with a PepMap100 RSLC, C18 3μm, 100Å, 75μm i.d. × 50cm column (Thermo Fischer Scientific) and the peptides were eluted with a linear gradient from 5-40% solvent B (A: 0.1% formic acid, B: 80% acetonitrile in 0.1% formic acid) in 90 minutes, at 300 nL/min flowrate.

For faster measurements, a Q-Exactive HF-X (Thermo Scientific, San Jose, CA, USA, Instrument Control Software version 2.9) was coupled to a UltiMate^TM^ 3000 RSLCnano system (Thermo Scientific, Germering, Germany) operated in capillary flow chromatography. Peptides were loaded onto a μ-Precolumn, 300 μm i.d × 5mm, C18 PepMap100, 5 μm, 100Å (Thermo Fisher Scientific) at a flow rate of t 50 μL/min for a min, loading solvent (2% ACN, 0.05% TFA). Then, the pre-column was switch online with a PepMap100 RSLC, C18 2μm, 100Å, 150μm i.d. × 15cm column (Thermo Fischer Scientific). The peptides were eluted with a linear gradient from 6-60% solvent B (A: 0.1% formic acid, B: 80% acetonitrile in 0.1% formic acid) in 34 minutes, at 1 μL/min flow rate. Mass spectrometer parameters settings in DDA, DIA and PRM modes on both instruments are described in the Supplementary methods.

### Spectral libraries generation

Proteome Discoverer 2.1.0.81 (Thermo Fischer Scientific) was used to search DDA raw files against Uniprot bacterial databases (databases are listed in Supplementary Methods). Peaklists were generated with the Spectrum Selector node (default parameters) of Proteome Discoverer and searched with using Mascot search engine version 2.5 (MatrixScience). Parameters were set for trypsin enzyme digestion specificity with two possible missed cleavages, methionine oxidation, asparagine and glutamine deamidation were set as variable modifications, and mass search tolerance were 10 ppm and 0.6 Da for MS and MS/MS respectively. Peptides were then validated at 1% FDR based on target/decoy search using Percolator software (33).

### Peptides selection and signal extraction in DIA analyses

For higher confidence and reproducibility in peptide identification, DIA signal extraction was performed on a selected part of the peptides identified in spectral libraries. Only peptides containing a maximum of 1 missed cleavage, having at least 8 amino acids in their sequence without any methionine and cysteine and identified in at least 6 Peptide Sequence Matches (PSM) were considered. The list of peptides from the 15 bacterial species was then searched with the Unipept software (34, 35) to delete peptide sequences also found in the human proteome, and to associate each peptide to the bacterial proteome it belongs to. Finally, for each bacterium separately, a list of potentially observable peptides was built and imported into Skyline 4.1.0.11796 (36, 37). Shuffle decoy peptides were added to allow further scoring. Spectral library was built in Skyline using the Proteome Discoverer results previously generated and using a cut-off score of 0.95. Retention time predictor was used considering the iRT peptides retention time values. Orbitrap resolving power was set at 30K at 200m/z with a high selectivity extraction. For each precursor (2+ or 3+), only 6 fragments (b or y) were automatically selected within 10 minutes around the predicted RT and their corresponding signal was extracted from the raw files. mProphet algorithm (38) was used within Skyline to score the peaks, considering the decoys and the second best peaks.

Only peaks with a Skyline dot product (dotP) > 0.75 and a *q*-value < 0.01 were considered as quantifiable and for each of them, the peptide areas (*i.e.* the sum of the area under the curve of the 6 most intense fragments) were normalized with the sum of the 10 iRT peptides. The non-quantifiable peaks received a value of 0. Finally, only the best intensity precursor of each peptide was kept. This list of peptides with the corresponding area values in each sample was used as an input for the machine learning algorithm.

### Peptidic signature generation by machine learning

We applied various machine learning models and several feature search approaches to identify a peptide signature. The peptide signature was composed of feature subsets found by three models having a good predictive performance: (i) Forward stepwise feature selection optimized by Matthew’s correlation coefficient criterion (39) using Naive Bayes classifier (40) with discretization, (ii) Forward stepwise feature selection optimized by Area Under the Curve (AUC) criterion using Bayesian Network classifier (41), and (iii) Forward stepwise feature selection combined with backward stepwise feature elimination optimized by Matthew’s correlation coefficient criterion using Hoeffding Tree classifier (42). All classifiers were evaluated by several subsampling procedures to assess the robustness of the models, including 10-fold cross validation, holdout and bootstrapping. Since stepwise feature selection tends to remove all correlated features, we retrieved those using Pearson and Spearman correlations, and having similar Information Gain score.

The obtained feature subsets of the three models were then merged, and manual curating was used to remove those observed in blank samples. The final curated signature was finally used to train a Bayesian Network prediction model. All new samples were analyzed using this predictive model.

### Peptidic signature validation and bacterial identification prediction

After PRM analysis using the Orbitrap Fusion or the Q-Exactive HF-X instrument, the Skyline software was used to extract the signal of the 82 peptides signature (*i.e*. the sum of the area under the curve of the 6 most intense fragments) in each sample. The peptides were considered as detected if they meet the following Skyline criteria: dotp > 0.85 and average mass error < 10 ppm, or dotp > 0.75 and average mass error < 3 ppm. For each analysis (inoculated urines or patient sample), the list of detected peptides was submitted to the Bayesian Network model trained in the previous section for prediction purposes.

### MALDI-TOF analysis

For all MALDI-TOF analyses, the standard procedure of the Enfant-Jésus hospital microbiology laboratory was used. Briefly, 1µL of urine was streaked on blood agar plate and 1µL on Mc Conkey agar plate (Oxoid). The plates were incubated 18 hours at 35°C. Isolated colonies with homogenous aspect were selected for MS analysis. The non-treated colonies were spotted on MALDI plate with HCCA matrix. MALDI-TOF MS analysis was performed on a Bruker Biotyper instrument using the Flux control version 3.4 (build 135) software and 7311 MSPs database.

### Experimental Design and Statistical Rationale

In order to obtain a high quality peptidic signature using machine learning algorithms, 9 high-level and 3 low-level inoculations replicates of each bacterial species were used. For the validation of the method in targeted proteomics, four different biological replicates of each bacterial inoculation in urine were monitored in two different analysis conditions. Finally, urine from 27 different patients were used to compare the method to conventional MALDI-TOF analysis. Prediction accuracies were reported.

## RESULTS

Our workflow for bacterial identification is composed of two steps: i) a training step which includes the acquisition of information on peptides expressed by the bacteria of interest using LC-MSMS in Data Independent Acquisition (DIA) mode followed by the generation of a short peptidic signature by machine learning models and ii) an identification step where the signature is monitored in unknown samples by PRM to obtain a bacterial identification through a prediction algorithm (Figure 1).

For the training step, in order to detect minor bacterial peptide signals in the human proteic background, we used DIA acquisition, on an Orbitrap Fusion instrument operating in nanoflow rate, because of its high sensitivity and its ability to provide a deep coverage of bacterial proteomes by acquisition of all peptides contained in the sample (43, 44). Indeed, in contrast to DDA which uses a full scan MS for the detection of peptide species, the DIA mode, by systematic acquisition of small size windows all along the mass range, improves the dynamic range and, thus, the sensitivity of the analysis. However, the simultaneous fragmentation of peptides inside this small window generates a complex spectrum which cannot be searched with conventional database search engines and needs to be deconvoluted to get peptide identifications.

### Acquisition of bacteria spectral libraries

One of the proposed approaches to extract information from the DIA complex spectra is to use spectral libraries previously acquired in DDA mode on the same type of sample and annotated with peptide/protein identifications through a protein database search (43). In our study, we have generated these spectral libraries from pure bacterial colonies in order to be as exhaustive as possible and cover a very wide range of bacterial tryptic peptides, and subsequently be able to extract this specific bacterial peptide information from the DIA complex spectra contaminated with human biological material.

We have generated spectral libraries for the 15 bacterial species of interest. To do so, each species was cultivated separately, proteins were extracted and digested with trypsin as described in the ‘experimental procedures’ section. The resulting peptides were fractionated by high-pH reversed phase chromatography. For each bacterial species, eight fractions were injected by LC-MS/MS in DDA mode and analyzed through a standard database search pipeline allowing the identification of 10686 to 29558 peptides at 1% FDR corresponding to 810 to 2438 protein groups (Supplementary Tables 1 and 2). As anticipated based on their genome size, gram-positive bacteria generated less protein identifications than the gram-negative. Indeed, there was a good correlation between the number of proteins identified in our study and the genome length (Pearson correlation coefficient *r* = 0.82) or the protein count predicted from genomic data (Pearson correlation coefficient *r* = 0.83) of all those 15 species. Thus, peptide fractionation combined to mass spectrometry analysis on a high resolution and high sensitivity instrument allowed us to cover 22.3 to 48.4 % of the Uniprot reference proteome of each of the 15 species (Supplementary Table 1). Then, the whole list of peptide identifications was refined to filter out: i) the peptides which may not to be reproducible from run to run (*i.e.* cysteine and methionine containing peptides, those containing trypsin missed cleavages, peptides shorter than eight amino-acids), and ii) the less abundant or less ionizable peptides (*i.e.* those having less than six Peptide Sequence Matches). Finally, we obtained a set of 31096 peptides which, according to their taxonomic affiliations, demonstrated a high redundancy across the 15 species (Supplementary Figure 2a).

This redundancy associated to our reproducibility filters showed that it is not possible to select from these data one or several specific peptides for each bacterial species that could would be further able to specifically sign for the presence of each distinct species in the urine. Indeed, not enough specific peptides are available when working which this large number of bacteria (*i.e.* 15) (Supplementary Figure 2b and 2c).

Thus, we aim to define a set of peptides that could be share by several species, but which, taken together, form a particular pattern for each bacterial species to be identified. To obtain this ‘peptidic signature’ our strategy was to use deep proteome coverage combined to machine learning algorithms to obtain this signature.

### Data Independent Analysis of artificially inoculated urine replicates

In order to define a peptidic signature of 15 bacterial species in the human urine background, we a have generated 12 artificial sample replicates, for each species of our selection, by inoculating urine from healthy volunteers with bacterial culture. Two concentration levels were used approximately set at 1×10^6^ CFU/mL (Colony Forming Unit per milliliter of urine) (n=9) (high level) and below 1×10^5^ CFU/mL (n=3) (low level) which corresponds to the threshold used by most of clinical laboratories for considering a UTI requiring an antibiotherapy. A total of 192 samples were produced, including ‘blank’ samples corresponding to non-inoculated urine as control. After protein extraction and short trypsin digestion, the resulting peptides were analyzed by LC-MS/MS in DIA mode. In comparison to DDA, DIA analyses enables a deep proteome coverage by reduction of the spectral dynamic range resulting in fewer missing values.

However, MS/MS spectra acquired in DIA mode are the sum of fragments generated by all precursor peptides selected in the same DIA window. It yields complex spectra where peptides sequences can be deduced by extraction of their specific fragments contained in the spectral libraries previously generated on pure bacterial colonies as described above. To do so, we have used the Skyline software (36, 37) and the list of 31096 selected peptides was used. An additional step of refinement was done to establish, for each species, the list of bacterial peptides to be searched for in the DIA runs. To this purpose, we used the Unipept software (34, 35) that enables to match peptide sequences with all matching taxa in UniProtKB databases. Starting from the non-redundant list of all peptides identified the bacteria, Unipept was used to confirm in which of the 15 bacterial species these could theoretically be found. Indeed, due to the stochastic effect of DDA used for library generation, it might be that some peptides belonging to several species had been sequenced by MS/MS in only a subset of them. The Unipept software also helped us to remove peptide sequences shared with the human proteome, hence generating high confidence lists of expressed peptides for each of the 15 species, free of potential human interfering compounds.

These lists and the corresponding spectra were added to Skyline for extracting DIA signals in the 12 replicates of each of the 15 inoculated samples. As retention time calibration peptides (iRT, Biognosys) were added in the DDA and DIA runs, predicted RT could be used for signal extraction in a small window of 10 minutes, thus limiting the probability for the software to select background peaks. A list of decoy peptides generated by Skyline were also extracted in the same conditions to ensure the calculation of a scoring *q*-value through the mProphet algorithm (38) included in Skyline.

Finally, the peptides were considered as detected if they met the following criteria: mProphet *q*-value < 0.01 and library dot product (dotp) > 0.75. After normalization of the peptide ions area values (i.e the sum of the 6 most intense fragments areas) by the sum of the iRT peptides area values and filtering for the best intensity precursor (when both doubly and triply-charged precursors were detected for the same peptide), the 15 final lists of peptides were combined into one and used as an input for machine learning algorithms to classify the bacteria and identify a short peptidic signature as described in the ‘experimental procedures section’. This computational method has been chosen for its ability to handle large datasets and to make prediction on them using accurate statistical models (45).

A list of 82 peptides was finally selected by our machine learning approach as the ‘peptidic signature’ allowing to discriminate the 15 most frequently found bacteria in UTI (Figure 2 and Supplementary Figure 3). This peptidic signature is composed of the peptide subsets found by three highly predictive models. Correlated peptides in the original dataset not retained in the models were added in the signature to improve the robustness of the final model, a Bayesian Network classifier. This model, trained on only high levels of concentration provided 100% classification accuracy on several *k*-fold cross-validations (*k* = 2, 5, 10) and was able to classify at 84% overall accuracy the low-level concentration samples (3 replicates per bacteria) corresponding to a concentration below the clinical threshold of 1×10^5^ CFU/mL.

**Figure 2.**
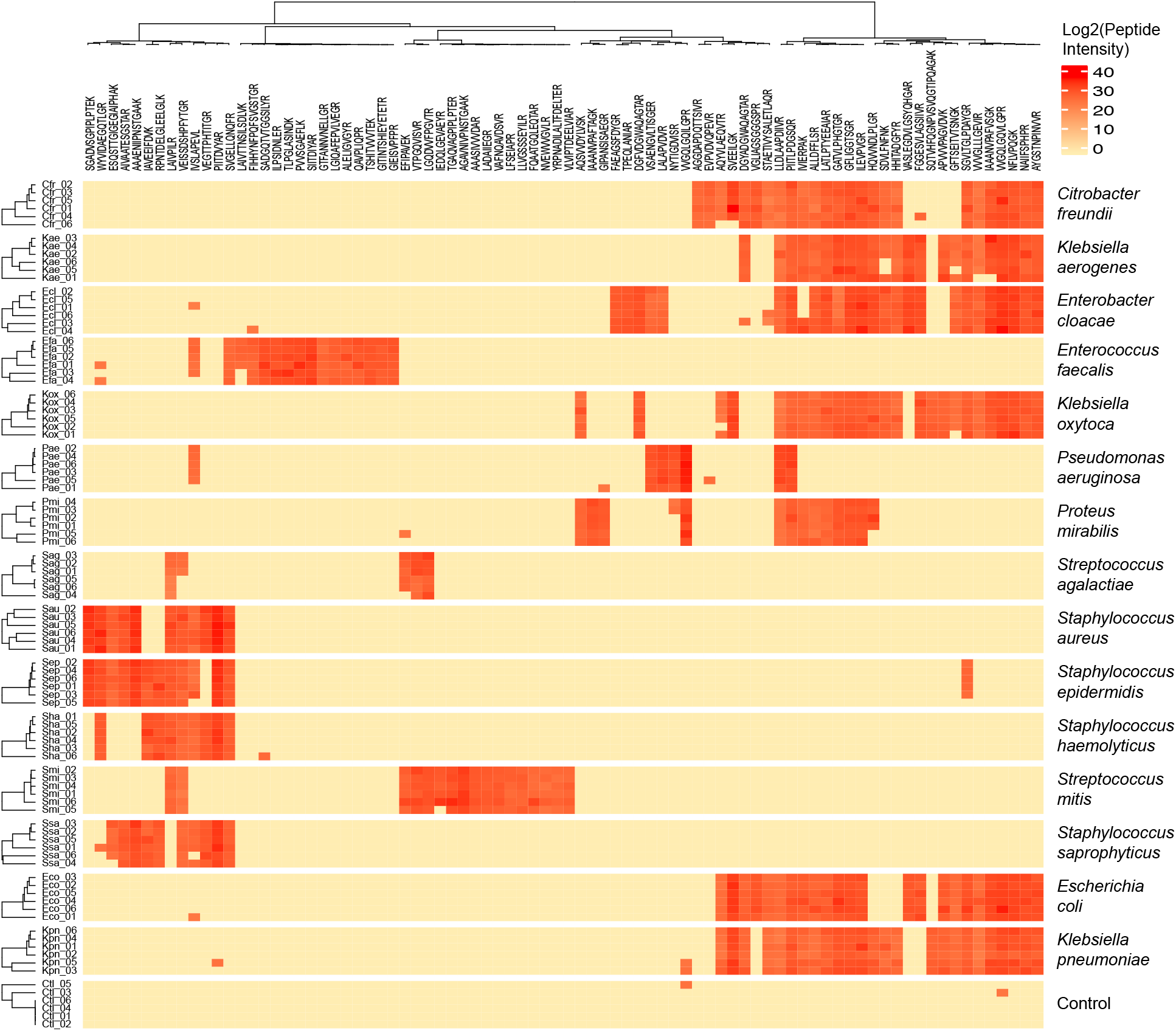
Heatmap of the peptidic signature corresponding to the 15 most frequently found bacteria in UTI. Intensity of each of the 82 peptides identified by the machine learning algorithm is represented for the six high-level concentration replicates of urine inoculation for each bacteria of interest. Data are presented with hierarchical clustering in rows and columns.

In the final signature, 5 to 26 peptides are observable for each bacterium. Even though closely related species, such as *Streptococcus epidermidis* and *Staphylococcus aureus*, or *Klebsiella pneumoniae* and *Escherichia coli*, share up to 75% of common peptides there are always a few peptides to distinguish them (4 and 7 peptides respectively in these two cases). For some very low concentration replicates, a few peptides, found high concentration replicates, were not detected. This loss affected the ability of the algorithm to predict the bacteria in only 15% of the tested low concentration replicates. Inversely, some false positive peptide detections were also observed, probably due to peak picking errors by Skyline in DIA runs, but they did not interfere with the bacterial prediction, assessing the robustness of the Bayesian Network model. As expected, most of the peptides composing the signature belong to relatively abundant proteins such as ribosomal proteins (*e.g.* 50S ribosomal protein L10, 30S ribosomal protein S5) or enzymes involved in amino acid metabolism (*e.g.* formate acetyltransferase) and glycolysis (*e.g.* GAPDH, pyruvate kinase) (46).

### Validation of the signature by targeted proteomics

Since the machine learning algorithm has identified a short list of peptides allowing the discrimination of the 15 bacteria of interest, this list can now be monitored by targeted proteomics which is known to give a better reproducibility of measurements and a better sensitivity in peptide detection and could thus improve the limit of detection of bacterial species in urine. The information on presence or absence of each of the 82 peptides of the signature is then given to the developed prediction model to obtain a probability of contamination. This step corresponds the Identification step of our pipeline (Figure 1). For this purpose, any type of mass spectrometer designed to perform targeted proteomics in Selected Reaction Monitoring (SRM) or Parallel Reaction Monitoring (PRM) modes can be used.

To validate our peptidic signature, we have initially used the Orbitrap Fusion Tribrid in PRM mode o which sample resulting from inoculated urines were injected. For this purpose, the four most frequently found bacteria in UTIs (*Escherichia coli, Streptococcus agalactiae, Enterococcus faecalis* and *Klebsiella pneumonia*) were inoculated at 5 different concentrations (from 2.56 × 10^4^ to 8.77 × 10^6^ CFU/mL) in urine from four different healthy volunteers (Supplementary Table 3). The samples were processed as described in the ‘experimental procedures’ section and analyzed with a 90 minutes gradient typically used in research laboratories. Only half of the volume of each sample was injected while the other half was kept for validation on other instrument types as further described.

The list of detected peptides for each replicate sample was used as an input for the prediction algorithm as new datasets to evaluate. The probabilities of bacterial identification are shown on Figure 3a and Supplementary Table 4. In 97% of the cases, the method was able to predict the correct species inoculated in the sample. In some cases, at low concentrations, the algorithm provided a false prediction, however in all cases except one, the prediction reports a “blank”. Finally, when looking at the data above the 1 × 10^5^ CFU/mL threshold commonly used by clinical labs, the accuracy in prediction reaches 100%.

**Figure 3.**
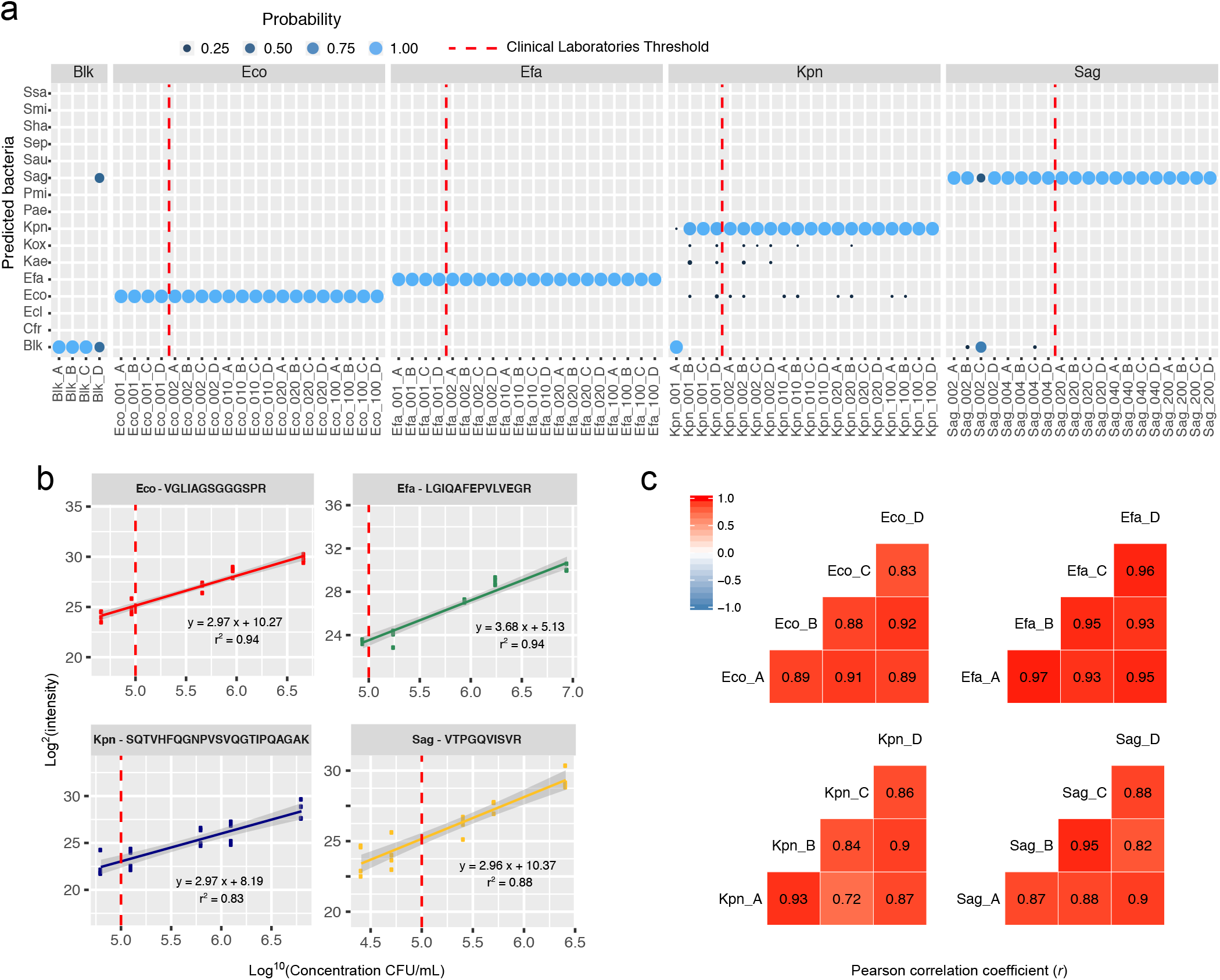
Accuracy, linearity and reproducibility of the ‘identification’ step of the method performed on the four most frequent bacteria in UTI. (a) Prediction reported by the algorithm after peptidic signature monitoring by PRM associated with its probability (light blue: high probability, dark blue: low probability) for five concentrations corresponding to five inoculation volumes (1, 2, 10, 20 and 100µL or 2, 4, 20, 40 and 200µL) of four bacteria (Eco, Efa, Kpn or Sag) in urine of four different healthy volunteers (A, B, C, and D), dotted red line corresponds to the commonly used clinical laboratories detection threshold of 1 × 10^5^ CFU/mL; (b) Linearity curves obtained for 4 peptides of the peptidic signature with the samples across the five tested concentrations, dotted red line corresponds to the commonly used clinical laboratories detection threshold of 1 × 10^5^ CFU/mL; (c) Pearson correlation coefficients between two of the four biological samples (*i.e* four different urines of healthy volunteers).

Using the intensities given by Skyline for each peptide, linearities curves have been plotted for each detected peptide of the signature (Figure 3b and Supplementary Figure 4a-d). The median of determination coefficient (R^2^) over the 5 concentrations was 0.841, suggesting that the method could be used for the quantification of bacterial contamination in biological samples. Since four biological replicates (i.e. bacterial inoculation in urines coming from four different volunteers) have been analyzed for each bacteria concentration, the reproducibility of the method was evaluated. Scatter plots and Pearson correlation factors were calculated from replicate to replicate (Figure 3c and Supplementary Figure 5a-d). For the same bacteria across various biological replicates, the Pearson correlation factors were 0.894 in average. This good reproducibility again suggests a possible use of the method for bacterial quantification in urine.

### Transfer of the signature on different instruments

To demonstrate that the initially designed signature using an Orbitrap Fusion instrument coupled to a nanoflow chromatography system is transferable to other instruments in others labs, we have analyzed the same inoculated urines of healthy volunteers (four different bacteria, five concentrations) in PRM mode on a Q-Exactive HF-X instrument coupled to capillary chromatography in PRM mode. Indeed, chromatography at higher flow rate (1 µL/min) improves the robustness of peptide separation and detection. To reduce the turnaround time between sample collection and bacterial identification as much as possible, the chromatographic gradient was also reduced between the Orbitrap Fusion and the Q-Exactive HF-X from 90 to 30 minutes. As for the Orbitrap Fusion data, the data collected from the Q-Exactive HF-X were analyzed using Skyline with the same validation criteria and the resulting list of detected peptides. Their intensity was used by the prediction algorithm (Figure 4a, Supplementary Figure 6 and Supplementary Table 4). Thus, in 94% of the cases, the method allowed the correct prediction of the bacteria initially inoculated in the samples. Errors were found only for some of the two lowest concentrations points and in all cases except one, the sample was predicted as blank. Again, when looking at the data above the clinical threshold of 1×10^5^ CFU/mL, the accuracy was significantly improved to reach 100%.

**Figure 4.**
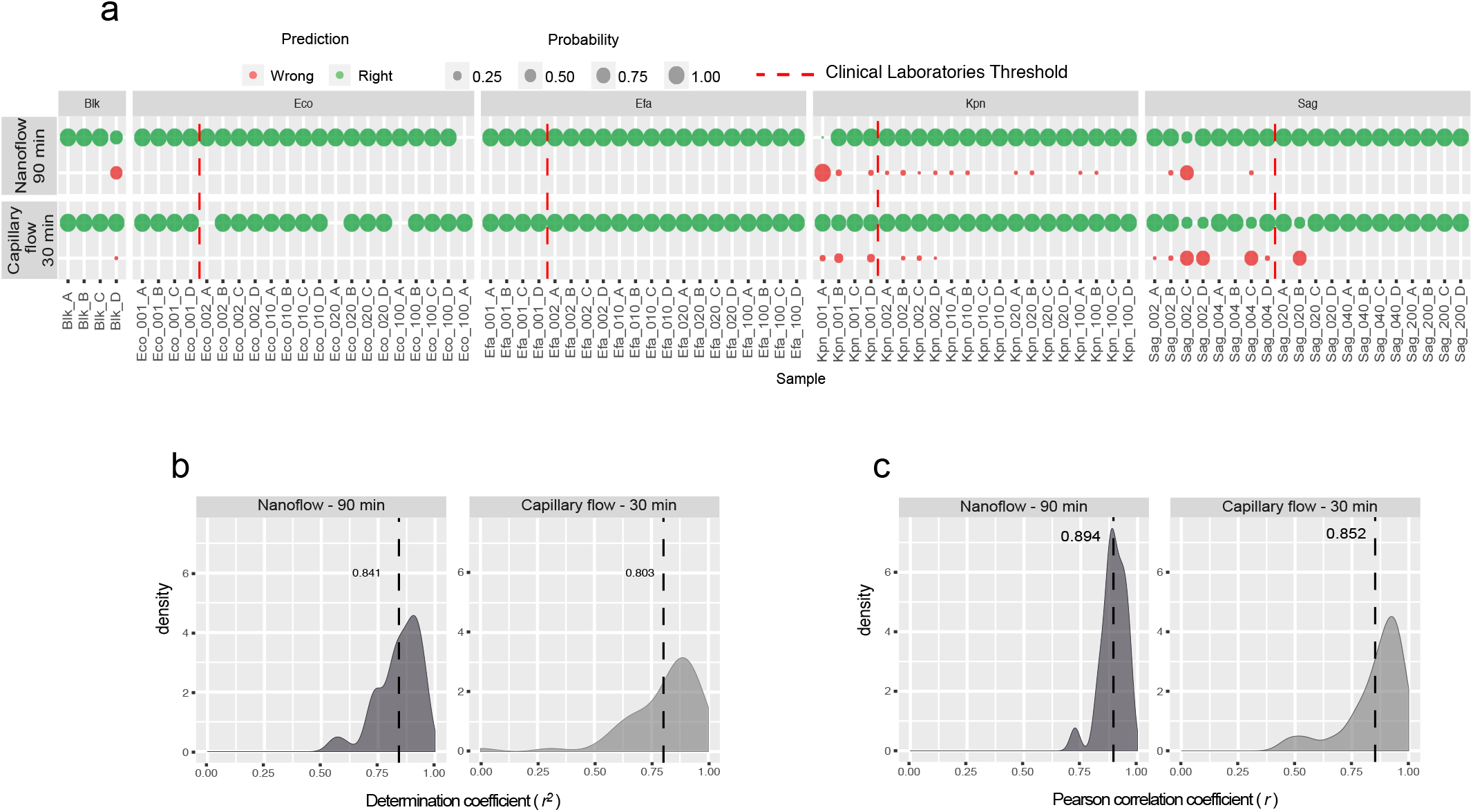
Accuracy, linearity and reproducibility of the ‘identification step’ of the method performed in two experimental conditions: 90 minutes gradient at nanoflow rate with PRM acquisition on an Orbitrap Fusion instrument or 30 minutes gradient at capillary flow rate with PRM acquisition on a Q Exactive HF-X instrument. (a) Right (green) or wrong (red) prediction reported by the algorithm after peptidic signature monitoring by PRM associated to its probability, for five concentrations corresponding to five inoculation volumes (1, 2, 10, 20 and 100µL or 2, 4, 20, 40 and 200µL) of four bacteria (Eco, Efa, Kpn or Sag) in urine of four different healthy volunteers (A, B, C, and D), dotted red line corresponds to the commonly used clinical laboratories detection threshold of 1e5 CFU/mL; (b) Distribution of the determination coefficients of the linearity curves obtained with the same samples across the five tested concentrations, the dotted line represents the average of all values; (c) Distribution of the Pearson correlation coefficients obtained by comparison of two biological replicates with the same samples, the dotted line represents the average of all values.

Linearities were also calculated by plotting the intensities of detected peptides across the five bacterial concentrations inoculated. The median of determination coefficients (R^2^) of all peptides was 0.803 (Figure 4b and Supplementary Figure 7a-d). In terms of reproducibility, four biological replicates were analyzed on the Q-Exactive HF-X. Scatter plots showed very good reproducibility since the Pearson correlation factors were 0.852 on average across the various biological replicates form the same bacterium at the same concentration (Figure 4c and Supplementary Figure 8a-d). However, lower Pearson correlation coefficient values were obtained for some peptides of *Streptococcus agalactiae.* This can be explained by the fact that this species has been inoculated at lower concentration than the others (Supplementary Table 3).

Again, the good results in terms of linearity and reproducibility obtained on the Q-Exactive HF-X instrument suggest its potential use for quantification of bacteria in urine.

### Validation on patient samples and comparison to MALDI-TOF MS

In order to validate our method in comparison to conventional MALDI-TOF analysis, samples were collected from 27 patients. Aliquots of the samples were analyzed using either our pipeline by monitoring the peptidic signature in PRM (nanoscale method) or with the standard MALDI-TOF method and the predictions of both methods were compared (Figure 5a and Supplementary Tables 4 and 5). Most of the analyzed urines were determined to be not infected (n=7) or infected by *E.coli* (n=9) with both methods, while 4 other samples contained 4 different bacteria among our 15 targeted species. For those 20 patients we found a correlation between MALDI-TOF and LC-MS in 95% of cases. In seven other cases, the LC-MS/MS reported the urine as ‘blank’ (not infected) while the MALDI-TOF reported an identification marked as ‘probable’ and limited at the genus level (*i.e.* without species mention). These results might be explain by a low level of contamination (below the level requiring anti-biotherapy) which prevented the LC-MS/MS method without culture to detect the signature peptides or by a contamination of the bacterial culture by non-pathogens, or infection of urines by species outside of our selection.

**Figure 5.**
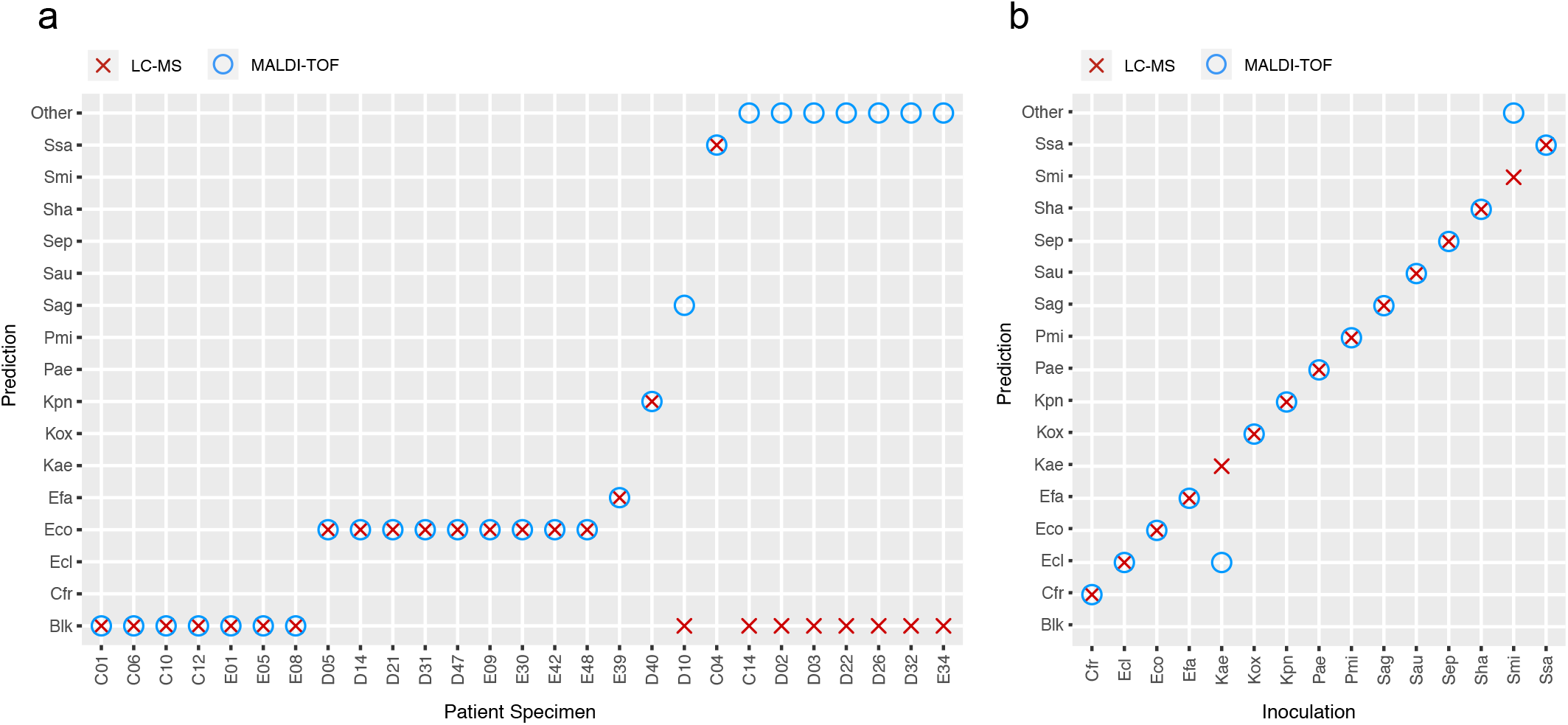
Comparison of the LC-MS method and the standard MALDI-TOF method. Prediction reported by the algorithm after peptidic signature monitoring by PRM without bacterial culture (red crosses) or by the MALDI Biotyper analysis after 24h hours bacterial culture (blue circles) on (a) 27 patients urine specimens or (b) 15 inoculations of bacterial species into urine from healthy volunteers.

Among the 15 bacterial species detectable by our method, many of them have a quite low frequency in UTI and not found in urines tested. In order to validate the detection of these species with our method in comparison with MALDI-TOF, we inoculated healthy urines with each of the 15 bacteria (Supplementary Table 3) and analyzed them with both pipelines. Our method found the correct inoculated bacterium in 100% of cases, while MALDI-TOF reported 2 errors (Figure 5b and Supplementary Tables 4 and 5). This lack of specificity in MALDI-TOF analysis might also explain some of the miscorrelations observed on patient samples.

## DISCUSSION

In this study, we developed a new strategy combining proteomics and machine learning for a fast, specific and accurate detection and identification of bacterial species present in urine without the need for time-consuming bacterial culture. We successfully applied our pipeline on the 15 bacterial species most commonly found in urine samples and obtained, in less than 4 hours, high rates of prediction accuracy, especially when looking above the quantitative threshold commonly used by clinical laboratories to consider a urine as infected and requiring anti-biotherapy.

This proof of concept could pave the way to the development of new peptide signatures for the analysis of other types of clinical specimens (bronchoalveolar lavage (BAL), stool, hemoculture…) (47–49), but also for the detection of foodborne or waterborne pathogens (50, 51), to reduce the turnaround time required to obtain a genus-and/or species-specific identification of microorganisms by classical or molecular microbiology methods or MALDI-TOF mass spectrometry. For some of these applications, without any culture, the sensitivity of the method might be too low to detect the bacteria but, it is expected that a short-term culture in a liquid medium might be enough to reach a detectable level (<1 × 10^3^ CFU/mL for certain peptides) without the isolation of colonies on a culture plate. In all cases, the high specificity of the method, due to a fine selection of the signature peptides, leads to a great improvement to what can be obtained with other standard methods such as MALDI-TOF mass spectrometry or 16S rRNA sequencing. This would be particularly valuable for the epidemiological surveillance of specific pathogens, instead of relying on expensive and time consuming whole genome sequencing (52, 53).

Moreover, the linearity and reproducibility of our method were evaluated and the obtained results suggest that the method could be used for quantification of bacterial cells in biological samples (for instance by addition of peptidic internal standards during the PRM monitoring). This would be particularly useful for urine specimens since a real infection needs to be distinguished from low level bacterial contaminants and this could serve to prevent the inappropriate use of antibiotics (54). This quantification is currently done by a plate counting of the bacterial culture which is a long and inaccurate process. As reported here, once a signature is created, it can be transferred to other laboratories or other targeted proteomics systems working in Parallel Reaction Monitoring (PRM) or Selected Reaction Monitoring (SRM) mode. Some of those instruments have already been approved as Medical Devices for other applications (55). Although the turnaround time to identify bacterial contaminants with this method is short (<4h), the non-parallelizable chromatographic time might limit its use in laboratories analyzing a high number of patient samples every day. However, we showed that this time can be reduced from 90 to 30 minutes and could be even more shortened with a better acquisition scheduling, some optimization of the LC gradient or use of high-throughput LC devices (56). Sample preparation time could also be automated and optimized, for instance, the trypsin digestion here done in one hour might be reduced to a couple of minutes as reported in the literature (57, 58). Finally, in a context where the emergence of resistant bacterial strains poses a global public health threat (59–61), the development of fast methods for bacterial typing becomes essential. Indeed, broad spectrum antimicrobials are commonly prescribed to patients before obtaining the results of the clinical microbiological analysis, a practice that might not be sufficient to control the infection, especially when one considers the risk posed by antibiotic-resistant species and their transmission in the hospital environment or the community (62). For instance, *Staphylococcus aureus* has developed resistances to many antimicrobial drugs including last resort antibiotics and expresses an arsenal of virulence factors (63–65). Or, in agriculture, the systematic use of antibiotics in farming leads to the selection of resistant bacteria that have been found in commercial food products (66). The accessibility with proteomics methods of the proteins involved in the resistance or virulence processes might constitute a challenge. However, several studies already reported the use of MALDI-TOF and LC-MS/MS to detect changes in the proteome of sensitive versus resistance strains (67–69). Thus, by including specific peptides belonging to proteins involved in resistance or virulence mechanisms in our peptidic signature, we could provide a measure of the risk associated to resistance or virulence and provide additional microbial information a physician could use to prescribe an appropriate antibiotic to a patient, thereby reducing the use of broad spectrum antibiotics.

Finally, we anticipate that the constant improvement in sensitivity, mass accuracy and acquisition speed of mass spectrometers will contribute in the future to improve the limit and precision of specific bacterial strains detection, making even more relevant the use of LC-MS/MS methods in microbiology.

## ABBREVIATIONS

DDA: Data Dependent Acquisition
DIA: Data Independent Acquisition
LC-MS/MS: Liquid Chromatography tandem Mass Spectrometry
MALDI-TOF: Matrix Assisted Laser Desorption Ionization – Time Of Flight
PRM: Parallel Reaction Monitoring
SRM: Selected Reaction Monitoring
UTI: Urinary tract infection(s)
Cfr: *Citrobacter freundii*
Ecl: *Enterobacter cloacae*
Eco: *Escherichia coli*
Efa: *Enterococcus faecalis*
Kae: *Klebsiella aerogenes*
Kox: *Klebsiella oxytoca*
Kpn: *Klebsiella pneumoniae*
Pae: *Pseudomonas aeruginosa*
Pmi: *Proteus mirabilis*
Sag: *Streptococcus agalactiae*
Sau: *Staphylococcus aureus*
Sep: *Staphylococcus epidermidis*
Sha: *Staphylococcus haemolyticus*
Smi: *Streptococcus mitis*
Ssa: Staphylococcus saprophyticus

## DATA AVAILABILITY

The raw mass spectrometry data have been deposited to the PASSEL repository with the identifier PASS01377 and will publicly accessible at http://www.peptideatlas.org/passel/ upon August 31^st^ 2019.

## ACKNOWLEDGMENTS

The authors are grateful to Benjamin Nehmé and Ève Bérubé for assistance in sample preparation, Geneviève Durand for MALDI-TOF analyses, and Anne Gonzalez de Peredo and Luc Bissonnette for critical reading of the manuscript.

