## SupplementaryMethods for "Fast and accurate bacterial species identification in biological samples using LC-MS/MS mass spectrometry and machine learning"

**Supplementary Methods**

*Bacterial Culture*

Bacterial strains were obtained from the Culture Collection of Centre de Recherche en Infectiologie of Laval University (CCRI, Quebec, Canada) registered as WDCM861 at the World Data Centre for Microorganisms. Their identification numbers as well as their culture conditions are listed below:

Supplementary Methods Table 1:
*Bacterial species and strains used*

| Abbreviation | Species | Strain | Culture condition | Gram-stain |
|---|---|---|---|---|
| Eco | *Escherichia coli* | CCRI-12923 | aerobe | negative |
| Efa | *Enterococcus faecalis* | CCRI-95$^T$ | aerobe | positive |
| Sag | *Streptococcus agalactiae* | CCRI-235$^T$ | air+5%$CO_2$ | positive |
| Kpn | *Klebsiella pneumoniae* | CCRI-563 | aerobe | negative |
| Pae | *Pseudomonas aeruginosa* | CCRI-691 | aerobe | negative |
| Sep | *Staphylococcus epidermidis* | CCRI-203 | aerobe | positive |
| Sau | *Staphylococcus aureus* | CCRI-9443 | aerobe | positive |
| Ssa | *Staphylococcus saprophyticus* | CCRI-223$^T$ | aerobe | positive |
| Smi | *Streptococcus mitis* | CCRI-259$^T$ | air+5%$CO_2$ | positive |
| Sha | *Staphylococcus haemolyticus* | CCRI-210$^T$ | aerobe | positive |
| Ecl | *Enterobacter cloacae* | CCRI-860 | aerobe | negative |
| Kox | *Klebsiella oxytoca* | CCRI-1109 | aerobe | negative |
| Cfr | *Citrobacter freundii* | CCRI-429$^T$ | aerobe | negative |
| Pmi | *Proteus mirabilis* | CCRI-675 | aerobe | negative |
| Kae | *Klebsiella aerogenes* | CCRI-446$^T$ | aerobe | negative |

T: denotes type strain. Identification of other strains was confirmed by gene sequencing and/or by MALDI-TOF

Each bacterial strain was plated on blood agar (TSA II 5% Sheep blood; Becton Dickinson) and incubated at 35°C overnight under aerobic conditions (or in air + 5% $CO_2$ for *S. agalactiae* and *S. mitis*). Bacterial colonies were then collected calibrated to a 0.5 MacFarland suspension in phosphate-buffered saline (PBS; 137mM NaCl, 6.4mM $Na_2HPO_4$, 2.7mM KCl, 0.88mM $KH_2PO_4$, pH 7.4). 30uL of this suspension were used to inoculate 3 mL of BHI medium (Brain Heart Infusion medium broth; Becton Dickinson) which was incubated overnight at 37°C under aerobic conditions with agitation (or 35°C in air + 5% $CO_2$ without agitation for *S. agalactiae* and *S. mitis*).
Finally, semi-log cultures were done by inoculation of 30µL of the previous culture in 3mL BHI medium and incubated in the same conditions. The cultures were stopped in exponential phase according to the previously known semi-log curve of each strain.

These cultures were used for spectral libraries generation or for urine inoculation. In parallel they were counted by incubation of 100uL of serial dilutions on Blood agar plates.

Supplementary Methods Table 2:
*Uniprot protein databases for DDA analyses*

| **Organism** | **Taxon ID** | **Uniprot Proteome ID** | **Number of entries** |
|---|---|---|---|
| *Citrobacter freundii* (Cfr) | 546 | UP000194801 | 5008 |
| *Enterobacter cloacae* (ECl) | 550 | UP000034104 | 4330 |
| *Escherichia coli* (Eco) | 83333 | UP000000625 | 4314 |
| *Klebsiella aerogenes* (Kae) | 1028307 | UP000008881 | 4909 |
| *Klebsiella oxytoca* (Kox) | 571 | UP000035545 | 6408 |
| *Klebsiella pneumoniae* (Kpn) | 272620 | UP000000265 | 5126 |
| *Pseudomonas aeruginosa* (Pae) | 208964 | UP000002438 | 5564 |
| *Proteus mirabilis* (Pmi) | 584 | UP000008319 | 3661 |
| *Enterococcus faecalis* (Efa) | 226185 | UP000001415 | 3240 |
| *Streptococcus agalactiae* (Sag) | 208435 | UP000000821 | 2105 |
| *Staphylococccus aureus* (Sau) | 93061 | UP000008816 | 2889 |
| *Staphylococcus epidermidis* (Sep) | 176280 | UP000001411 | 2461 |
| *Staphylococcus haemolyticus* (Sha) | 279808 | UP000000543 | 2640 |
| *Streptococcus mitis* (Smi) | 28047 | UP000027992 | 1983 |
| *Staphylococcus saprophyticus* (Ssa) | 342451 | UP000006371 | 2404 |
| *Homo sapiens* | 9606 | UP000005640 | 73928 |

Supplementary Methods Table 3:
*Mass Spectrometry parameters*

| | **DDA** | **DIA** | **PRM** | **PRM** |
|---|---|---|---|---|
| **Instrument** | Orbitrap Fusion | Orbitrap Fusion | Orbitrap Fusion | Q-Exactive H-FX |
| Method duration | 120 min | 120 min | 120 min | 50 min |
| Polarity | Positive | Positive | Positive | Positive |
| ***MS parameters*** | | | | |
| Analyzer | Orbitrap | Orbitrap | | |
| Resolution | 120K | 60K | | |

| | | | | |
|---|---|---|---|---|
| Mass Range | 350-1800 | 400-1000 | | |
| AGC | 4E+05 | 4E+05 | | |
| Max Injection time (ms) | 50 | 50 | | |
| ***MSMS parameters*** | | | | |
| Isolation window | 1.6 | 10 (40 windows, 400-800) | 0.7 | 0.7 |
| Activation type | HCD | HCD | HCD | HCD |
| Collision energy | 35% | 35% | 35% | 28 |
| Analyzer | Ion Trap | Orbitrap | Orbitrap | Orbitrap |
| Resolution or Scan rate | Rapid | 30K | 30K | 15K |
| AGC | 1E+04 | 4E+05 | 5E+04 | 1E+05 |
| Max Injection time (ms) | 50 | 70 | 120 | 100 |
| Data Dependent MS2 parameters | Most intense precursor with intensity greater than 5000, Top speed 3s, Dynamic exclusion 20s with 10ppm tolerance. | | | |
| Internal calibration | Lock mass 445.12003 | Lock mass 445.12003 | Lock mass 445.12003 | Lock mass 445.12003 |
