## SupplementaryMaterial for "Fast and accurate bacterial species identification in biological samples using LC-MS/MS mass spectrometry and machine learning"

Supplementary Figure 1. **Statistics of Urine Specimen analyses at the microbiology laboratory of Enfant-Jesus Hospital in Quebec City, Canada.** The pie chart represents, for infected specimens, the proportion of each bacterial species identified by MALDI-TOF analysis using the standard procedure described in the Methods section over four months (January to April 2017).

Supplementary Figure 2. **Sequence Redundancy between the 15 bacterial species of interest.** (a) Heatmap represented the list of 31893 peptides sequences selected after DDA analysis of each pure species. Each sequence found in a species is indicated in dark blue. Hierarchical clustering was performed in rows and columns. (b) Using the same list of peptide sequences, the graph indicates how many peptides are specific to only one species when comparing 2 or more species (in frequency order in UTIs). (c) The bar plot represents the repartition of specific peptides when comparing 15 bacterial species.

Supplementary Figure 3. **Heatmap of the peptidic signature corresponding to the 15 most frequently found bacteria in UTI.** Intensity of each of the 82 peptides identified by the Machine Learning algorithm is represented for the all low-level and high-level concentration replicates of urine inoculation for each bacteria of interest. Data are presented with a hierarchical clustering in rows and columns.

Supplementary Figure 4. **Linearity of the 'identification step' of the method performed with a 90 minutes gradient on an Orbitrap Fusion instrument.** Linearity curves were plotted for five concentrations corresponding to five inoculation volumes (1, 2, 10, 20 and 100μL or 2, 4, 20, 40 and 200μL) in urine of four different healthy volunteers (A, B, C, and D) for peptides of four bacteria: (a) *Escherichia coli*, (b) *Enterococcus faecalis*, (c) *Klebsiella pneumoniae*, (d) *Streptococcus agalactiae*. Dotted red line corresponds to the commonly used clinical laboratories detection threshold of 1e5 CFU/mL.

Supplementary Figure 5. **Reproducibility the 'identification step' of the method performed with a 90 minutes gradient on an Orbitrap Fusion instrument.** Scatter plots and Pearson correlation coefficient are presented for four biological replicates corresponding to all inoculation volumes in urine of four different healthy volunteers (A, B, C, and D) for four bacteria: (a) *Escherichia coli*, (b) *Enterococcus faecalis*, (c) *Klebsiella pneumoniae*, (d) *Streptococcus agalactiae*.

Supplementary Figure 6. **Accuracy of the 'identification step' of the method performed with a 30 minutes gradient on a Q-Exactive HF-X instrument.** Predictions reported by the algorithm after peptidic signature monitoring by PRM associated with its probability (light blue : high probability, dark blue : low probability) for five concentrations corresponding to five inoculation volumes (1, 2, 10, 20 and 100μL or 2, 4, 20, 40 and 200μL) of four bacteria (Eco, Efa, Kpn or Sag) in urine of four different healthy volunteers (A, B, C, and D), dotted red line corresponds to the commonly used clinical laboratories detection threshold of 1e5 CFU/mL.

Supplementary Figure 7. **Linearity of the 'identification step' of the method performed with a 30 minutes gradient on a Q-Exactive HF-X instrument.** Linearity curves were plotted for five concentrations corresponding to five inoculation volumes (1, 2, 10, 20 and 100μL or 2, 4, 20, 40 and 200μL) in urine of four different healthy volunteers (A, B, C, and D) for peptides of four bacteria: (a) *Escherichia coli*, (b) *Enterococcus faecalis*, (c) *Klebsiella pneumoniae*, (d) *Streptococcus agalactiae*. Dotted red line corresponds to the commonly used clinical laboratories detection threshold of 1e5 CFU/mL.

Supplementary Figure 8. **Reproducibility the 'identification step' of the method performed with a 30 minutes gradient on a Q-Exactive HF-X instrument.** Scatter plots and Pearson correlation coefficient are presented for four biological replicates corresponding to all inoculation volumes in urine of four different healthy volunteers (A, B, C, and D) for four bacteria: (a) *Escherichia coli*, (b) *Enterococcus faecalis*, (c) *Klebsiella pneumoniae*, (d) *Streptococcus agalactiae*.

Supplementary Table 1. **Summary of identification results obtained by LC-MSMS analysis of each pure bacterial species in DDA mode correlated with genomic data.** The table presents the number of proteins and peptides identified after database search of the MS data in Uniprot. The proteome coverage was calculated using the percentage of proteins identified regarding the number of entries in the database. The results were compared to the Total genomic length and Predicted protein count of each species found in NCBI genome database.

Supplementary Table 2. **Proteome Discoverer 2.1 identification results for each pure bacterial species analyzed by LC-MSMS in DDA mode**. Peptide export obtained after Mascot search and Percolator validation. Only high confidence peptides were kept. Each tab corresponds to one of the 15 bacteria of interest.

Supplementary Table 3. **Count of bacterial cultures reported to urine inoculation.** Bacterial cultures were counted as described in the Methods section and the values obtained were used to determine the bacterial concentration in urine inoculates.

Supplementary Table 4. **Bacterial identification reported by MALDI-TOF and LC-MSMS analyses.** For samples analyzed with both methods, the "MALDI-TOF" column shows the results obtained with the standard procedure at Enfant-Jesus Hospital of Quebec City, Canada, the "LC-MS" column shows the prediction given by the algorithm with our method.

Supplementary figure 1

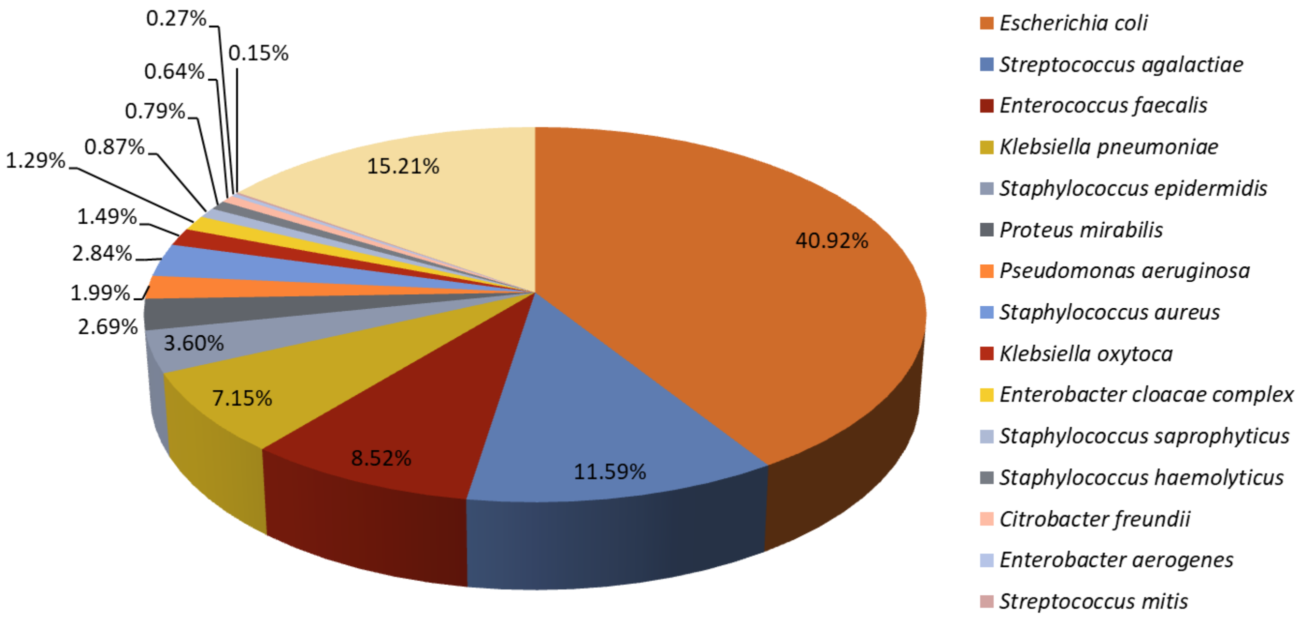

Supplementary Figure 2

a

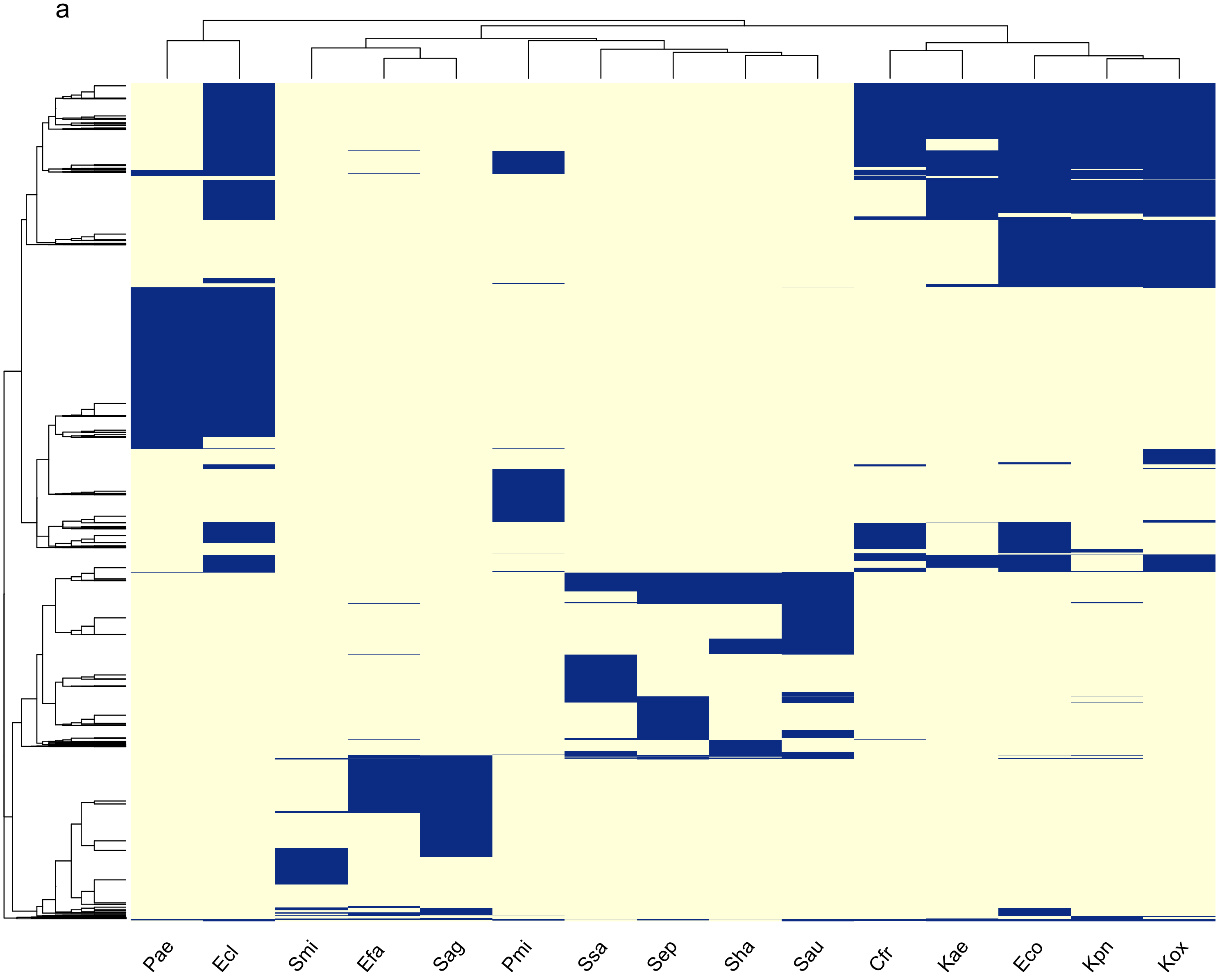


b

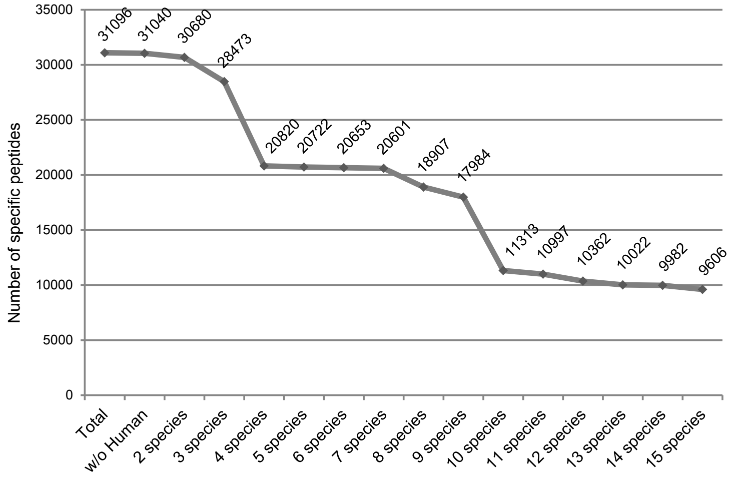


c

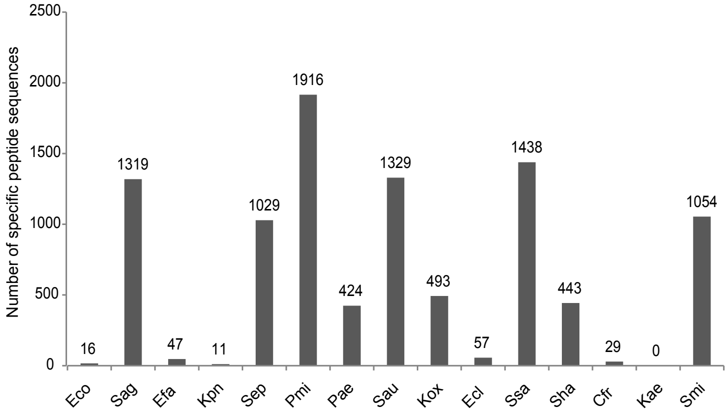

### Supplementary Figure 3

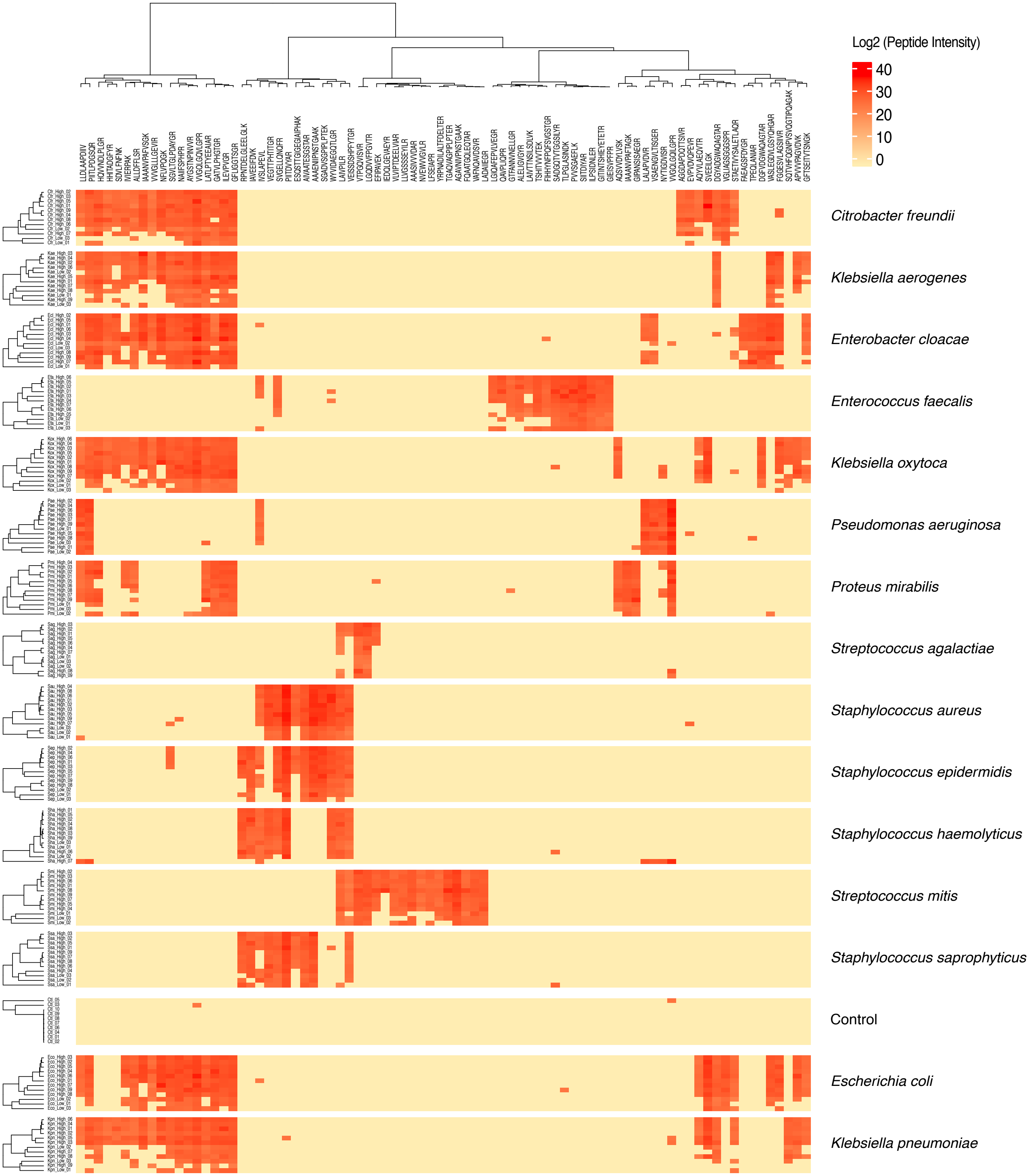

Supplementary Figure 4

a

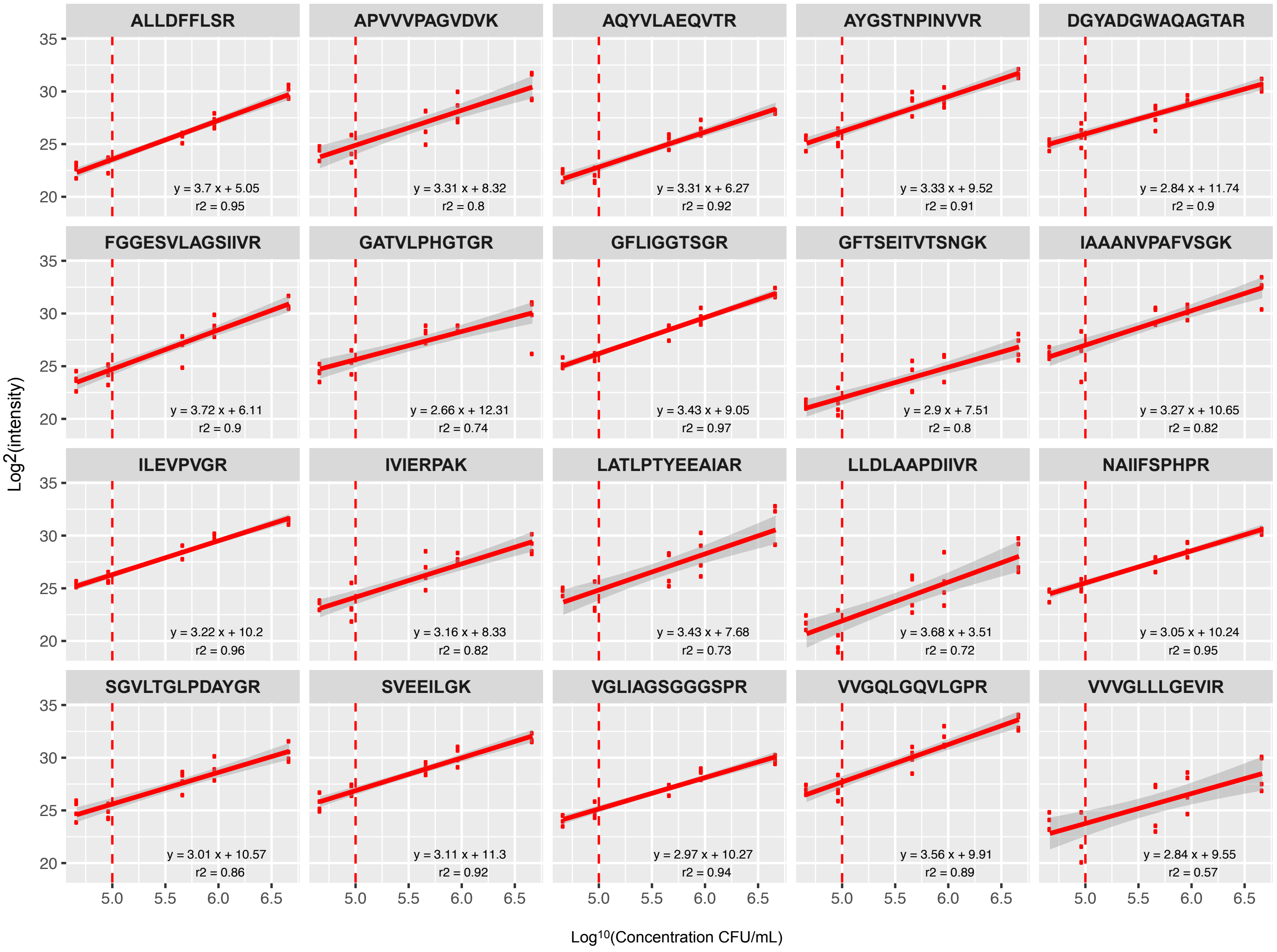

Supplementary Figure 4

b

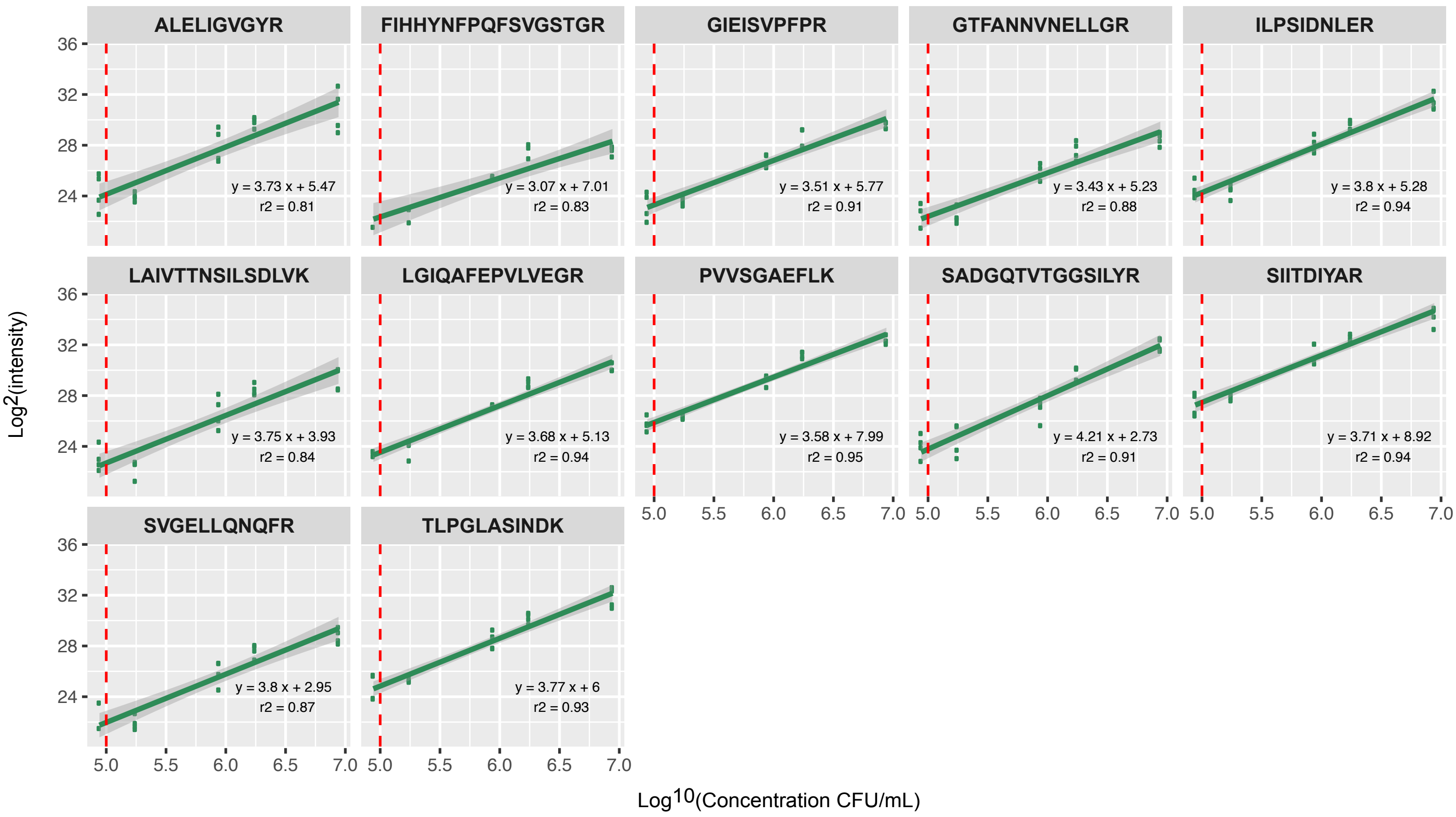

Supplementary Figure 4

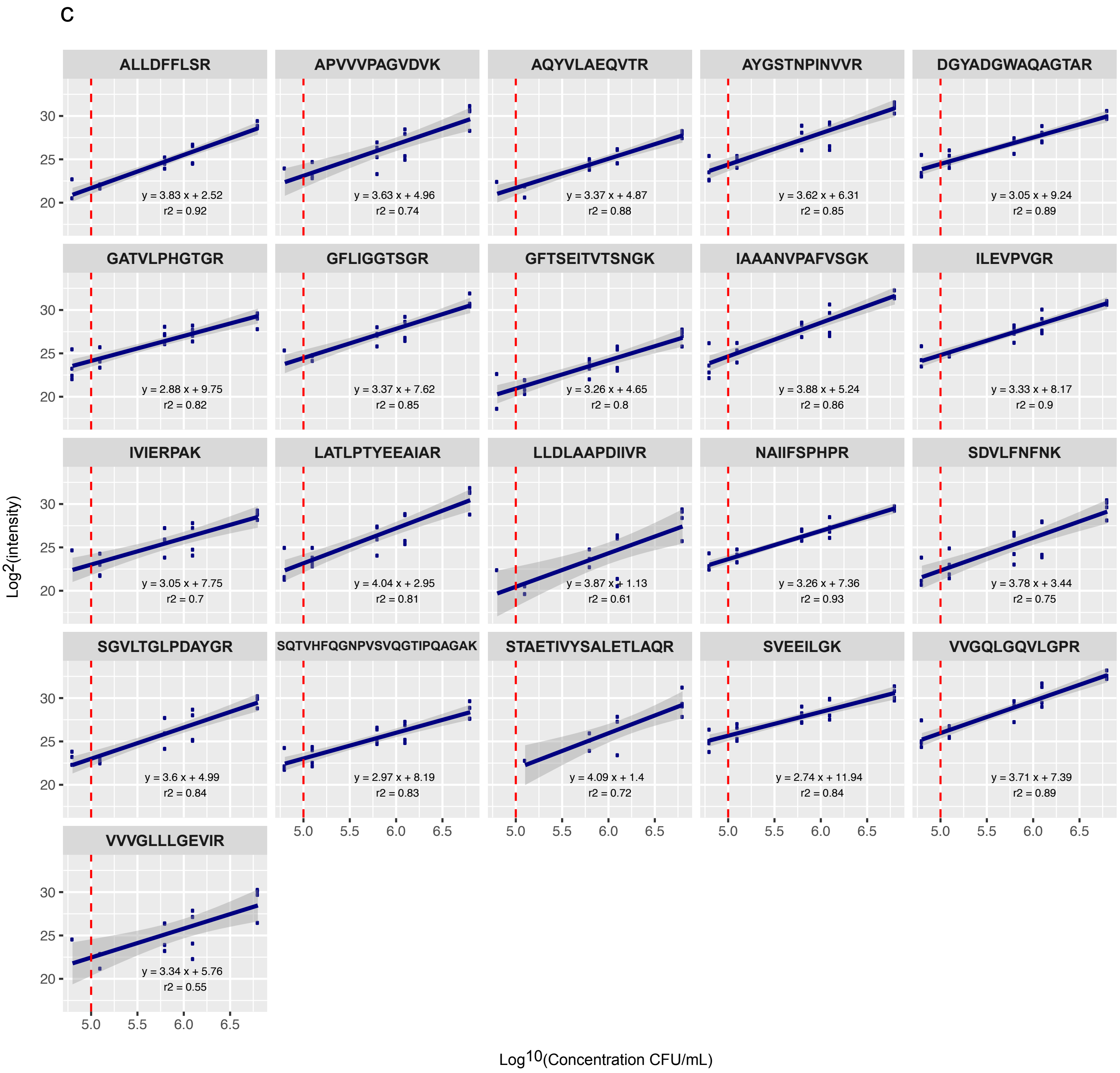

### Supplementary Figure 4

d

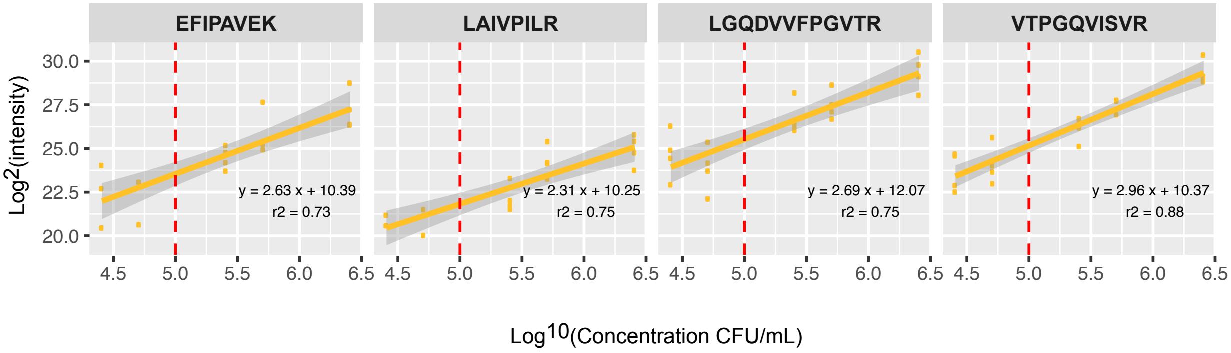

Supplementary Fig5

a

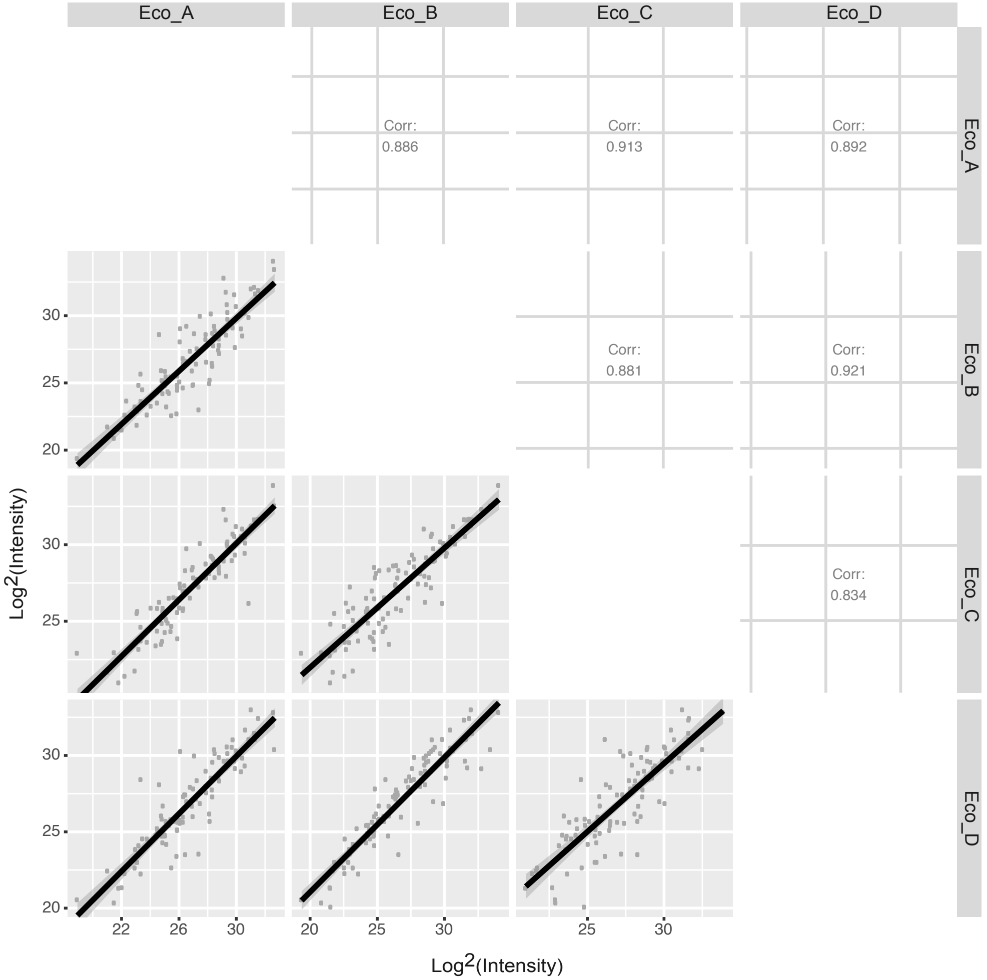


b

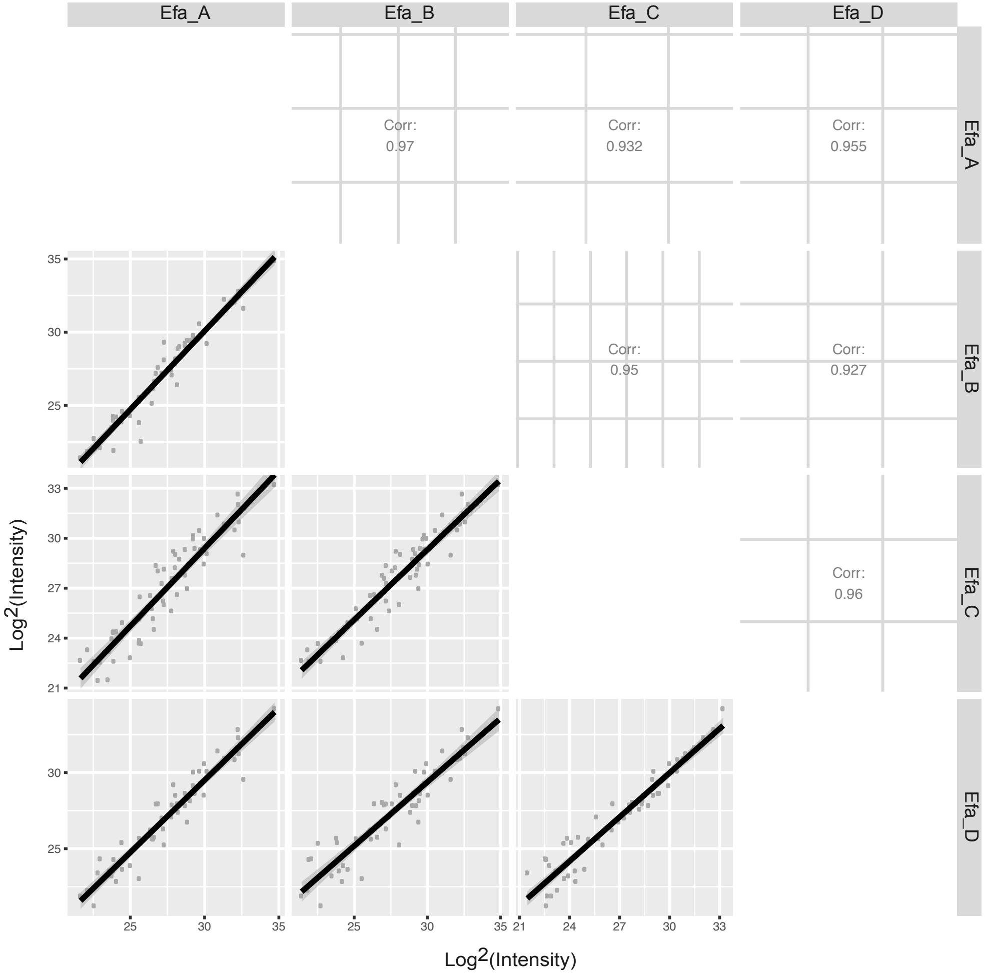

### Supplementary Figure 5

c

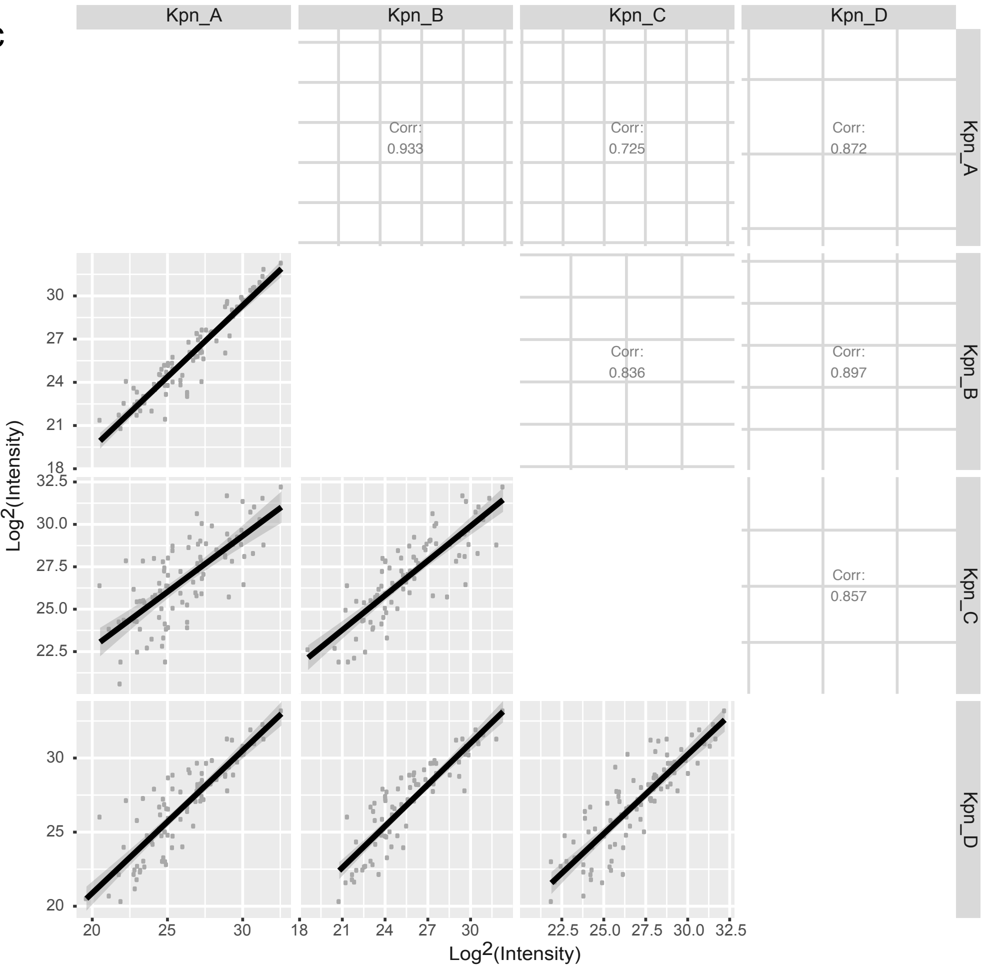


d

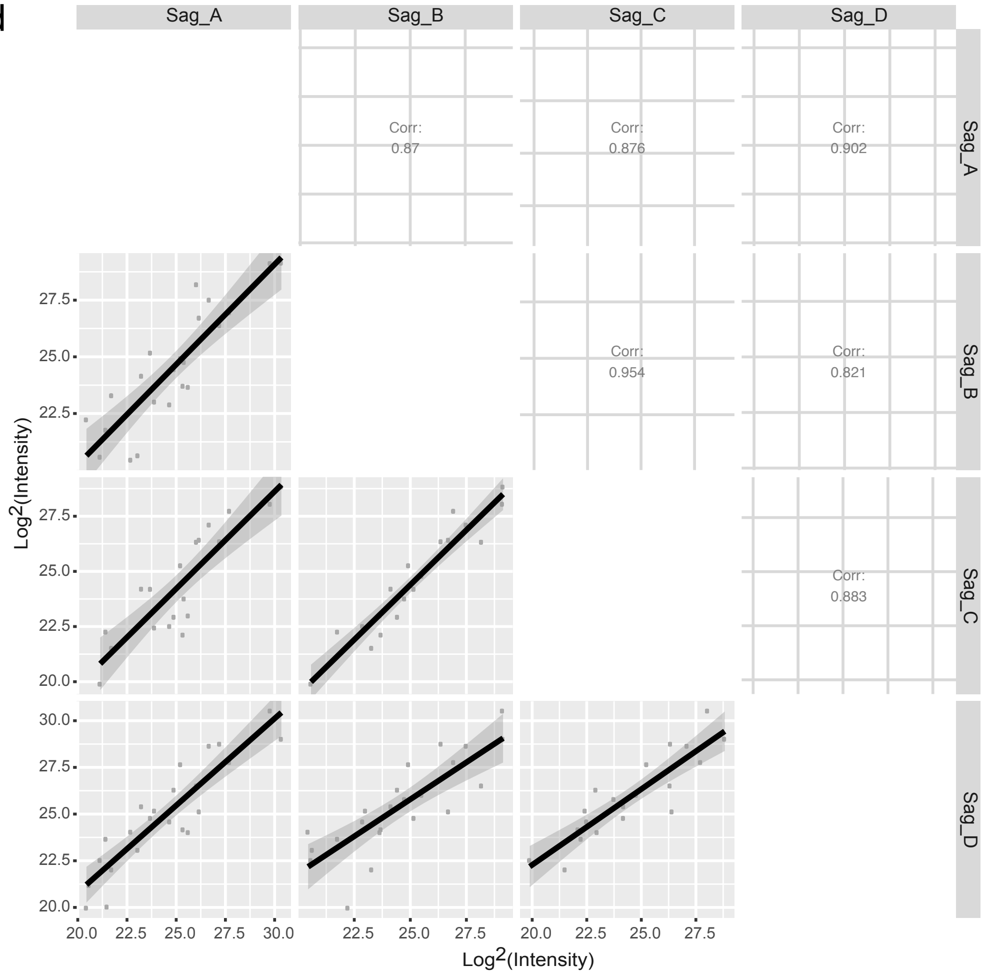

### Supplementary Figure 6

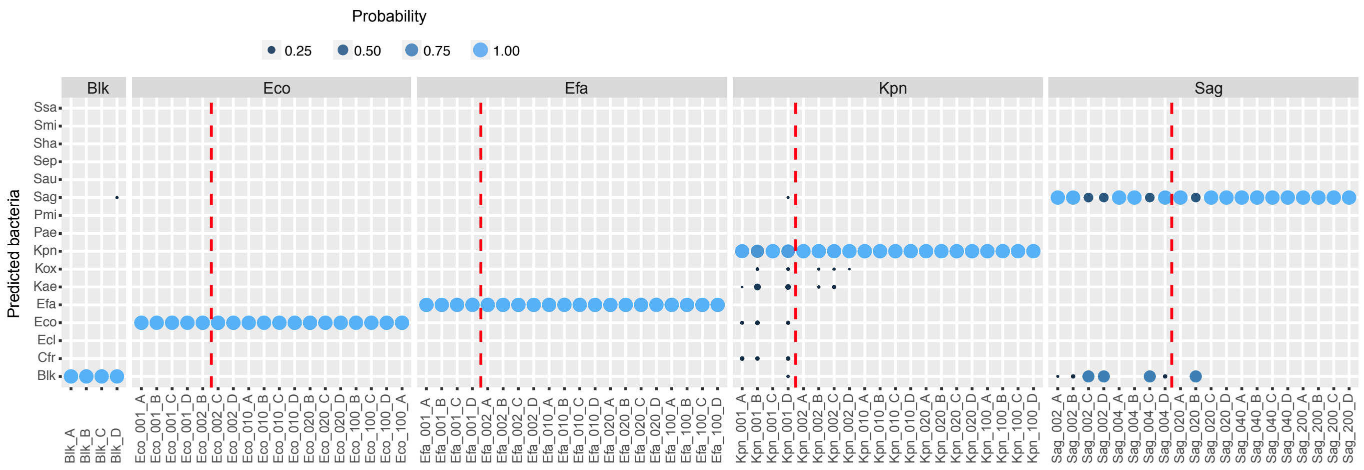

Supplementary Figure 7

a

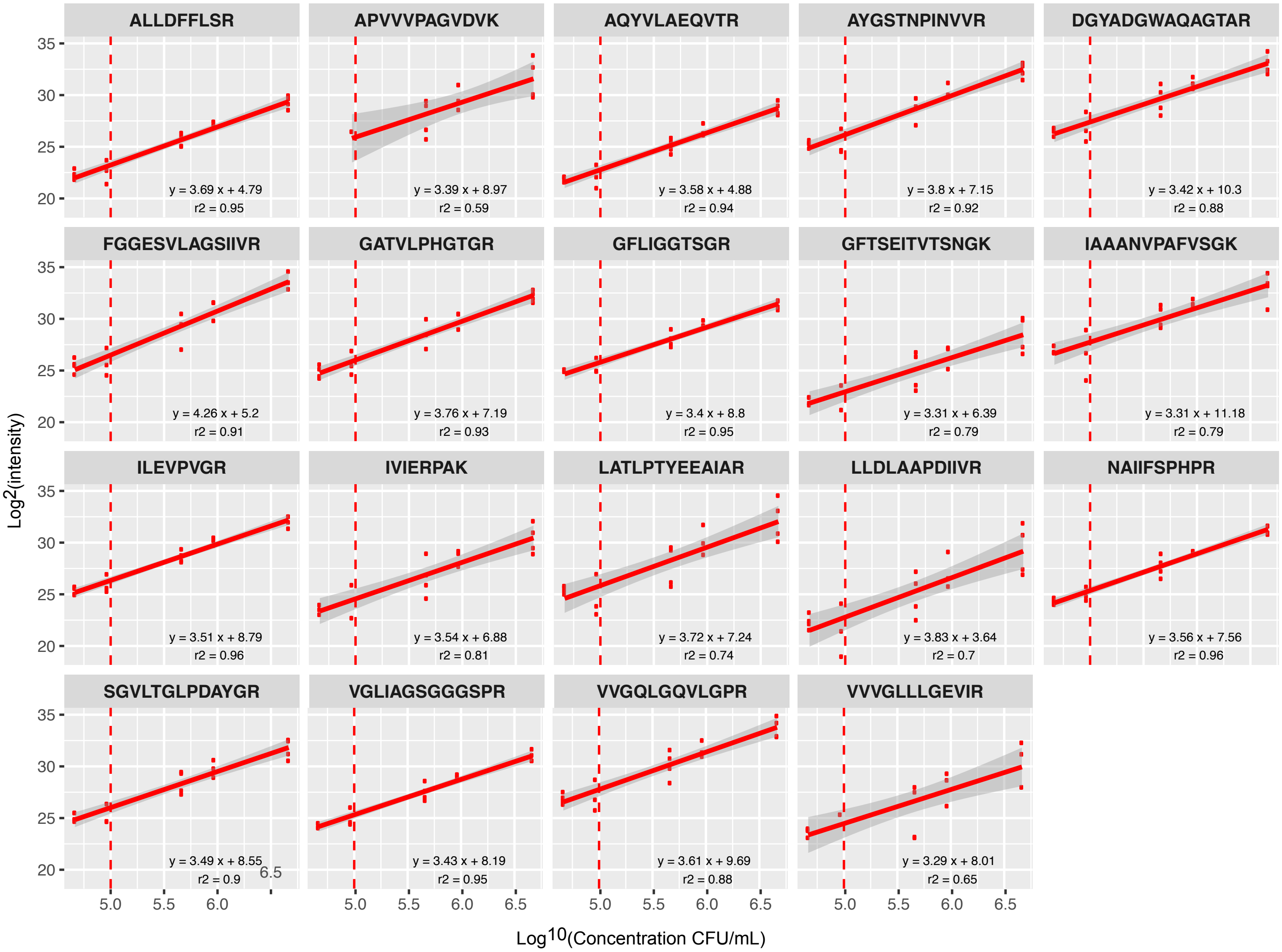

Supplementary Figure 7

b

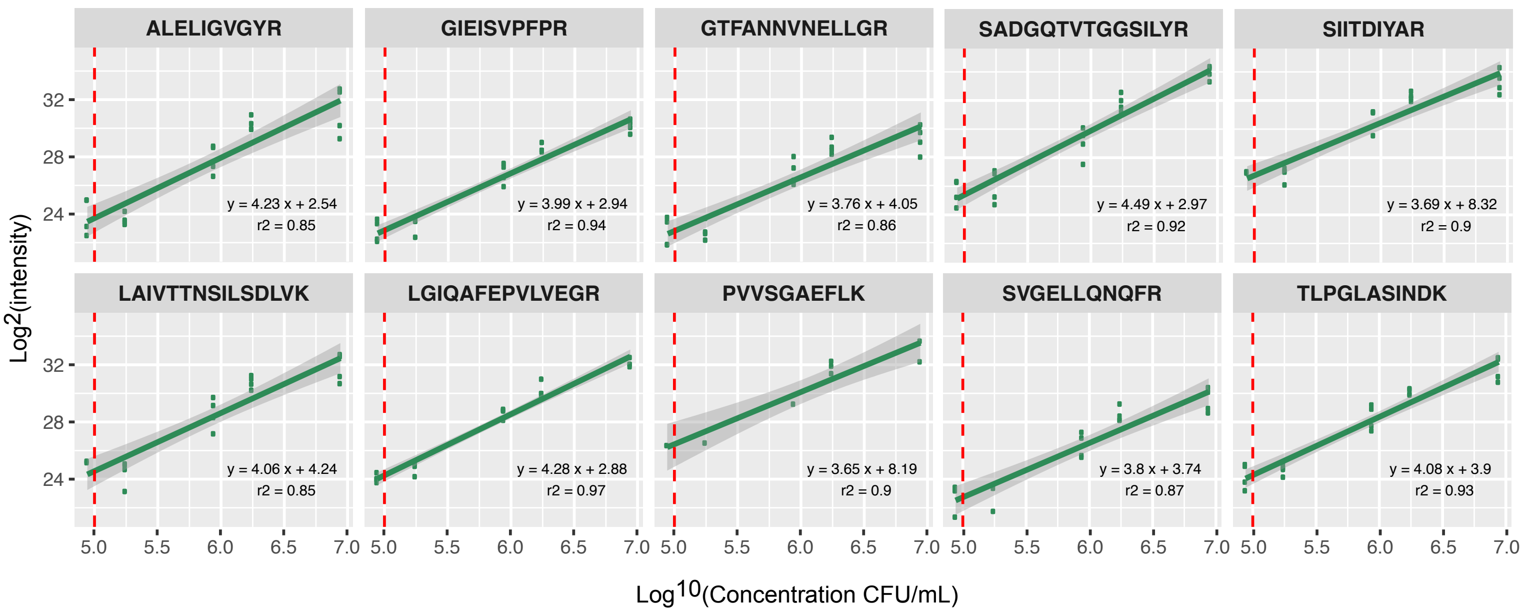

Supplementary Figure 7

c

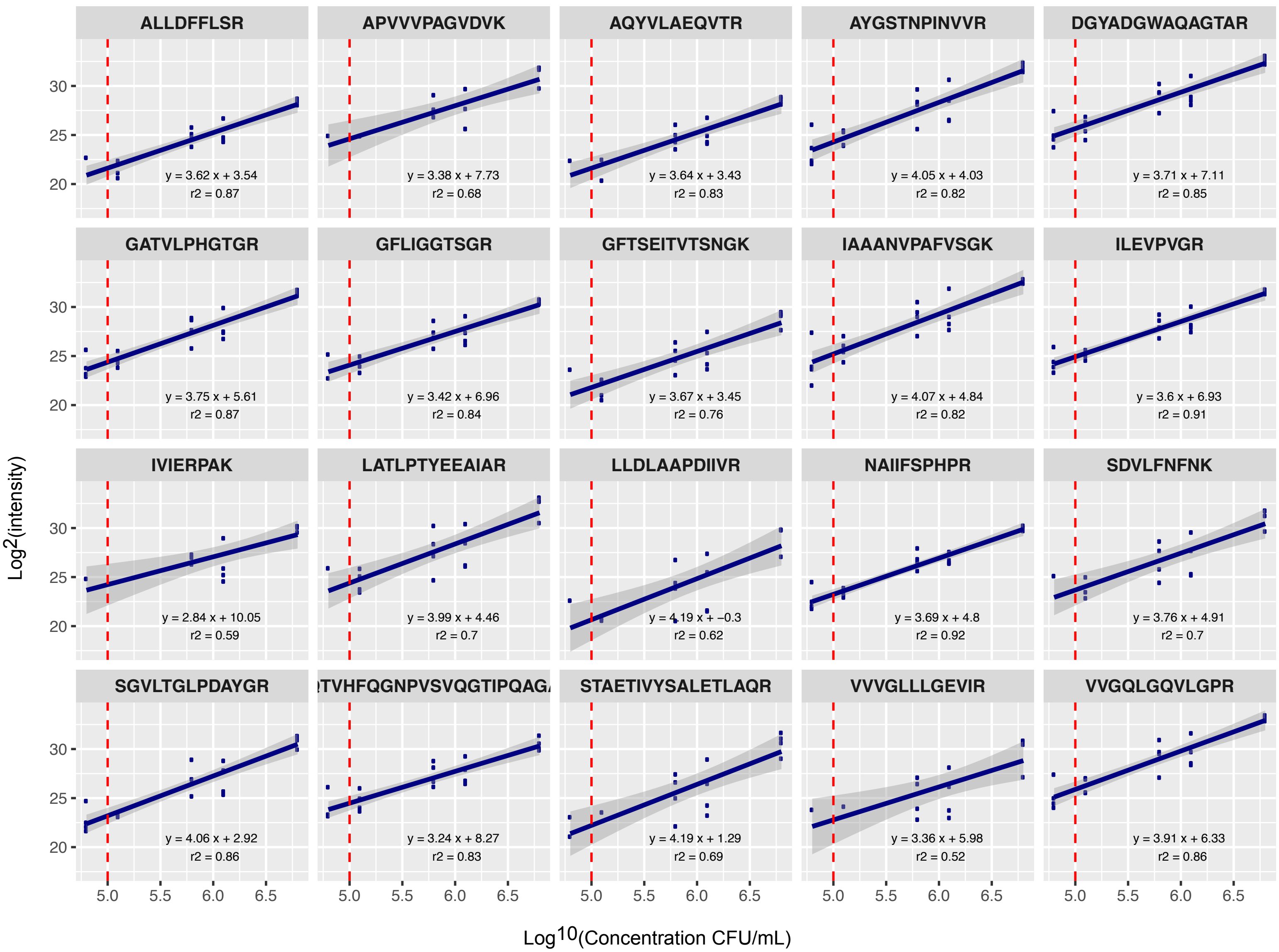

Supplementary Figure 7

d

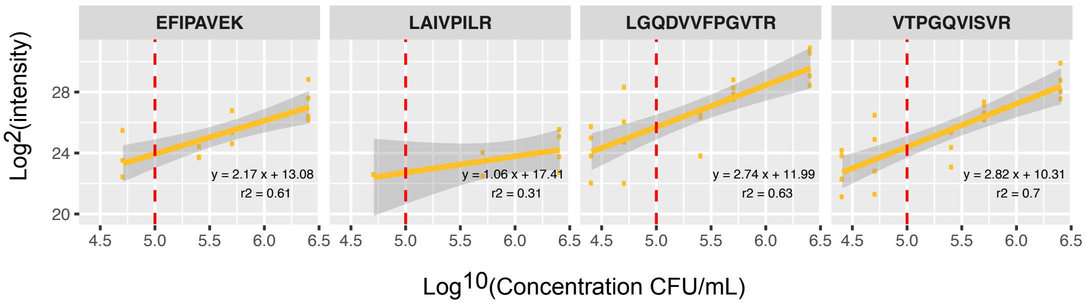

Supplementary Figure 8

a

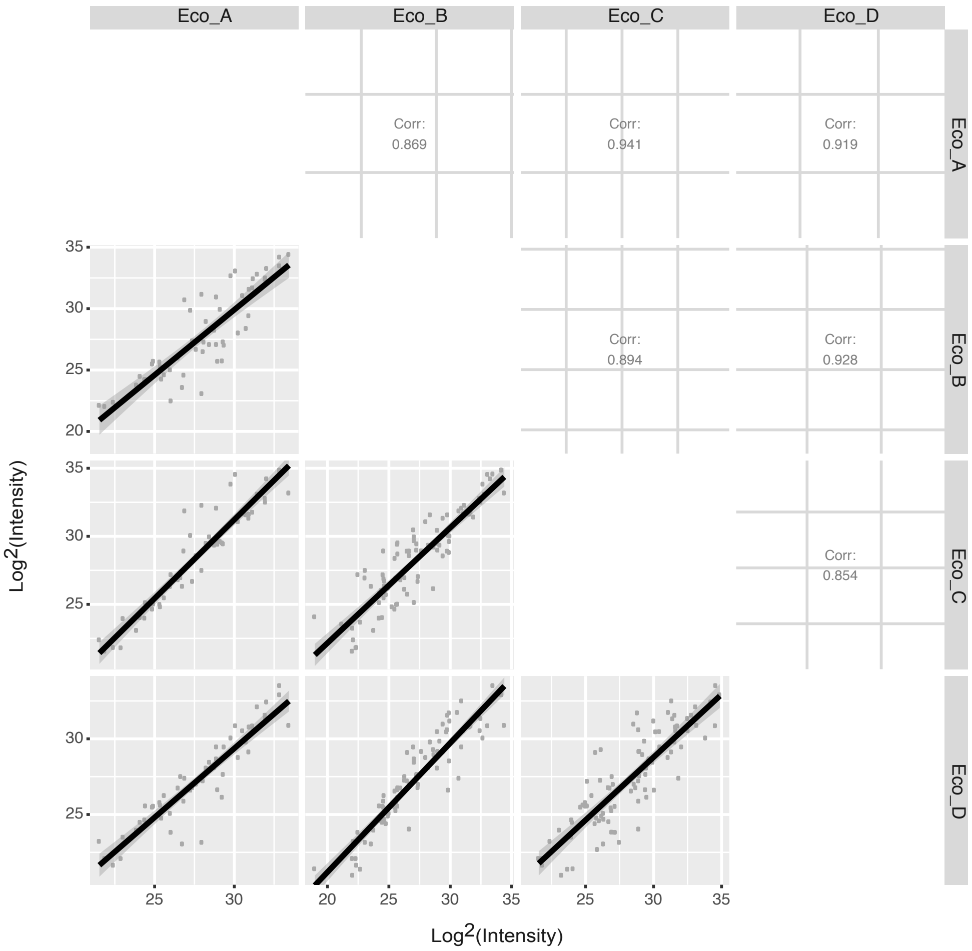


b

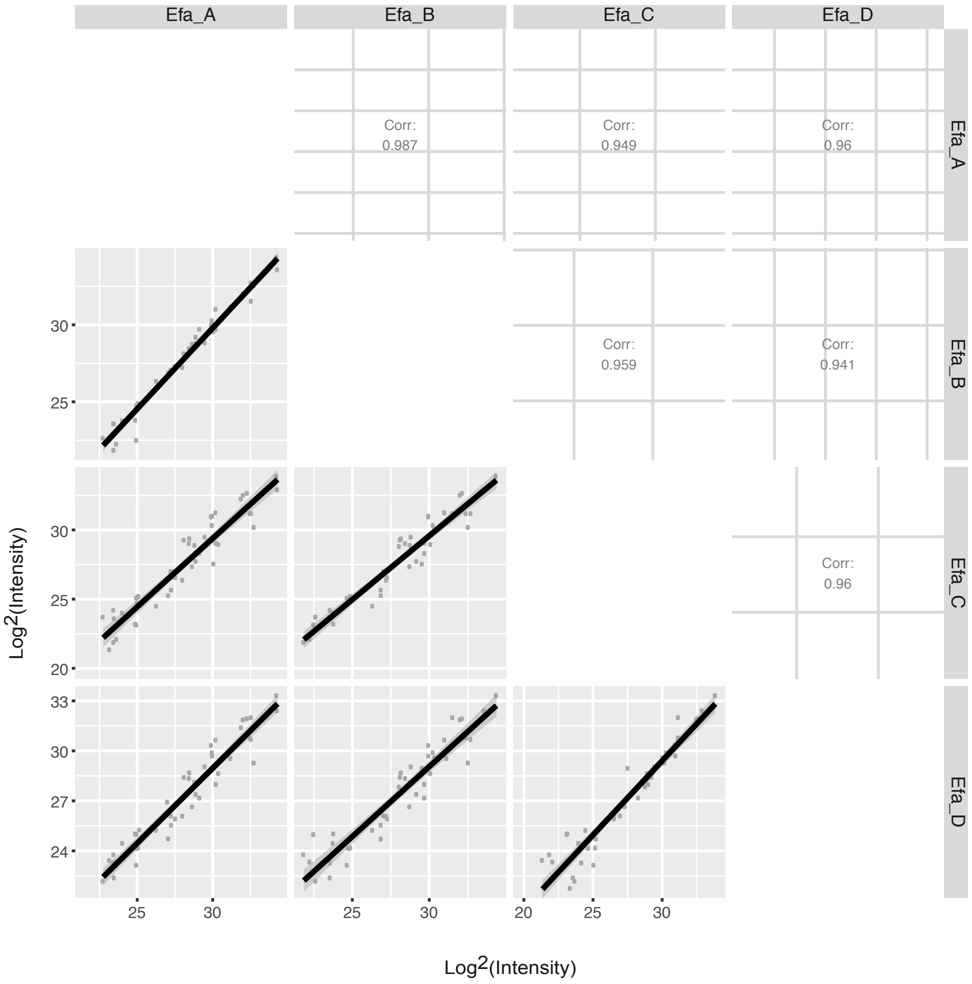

Supplementary Figure 8

c

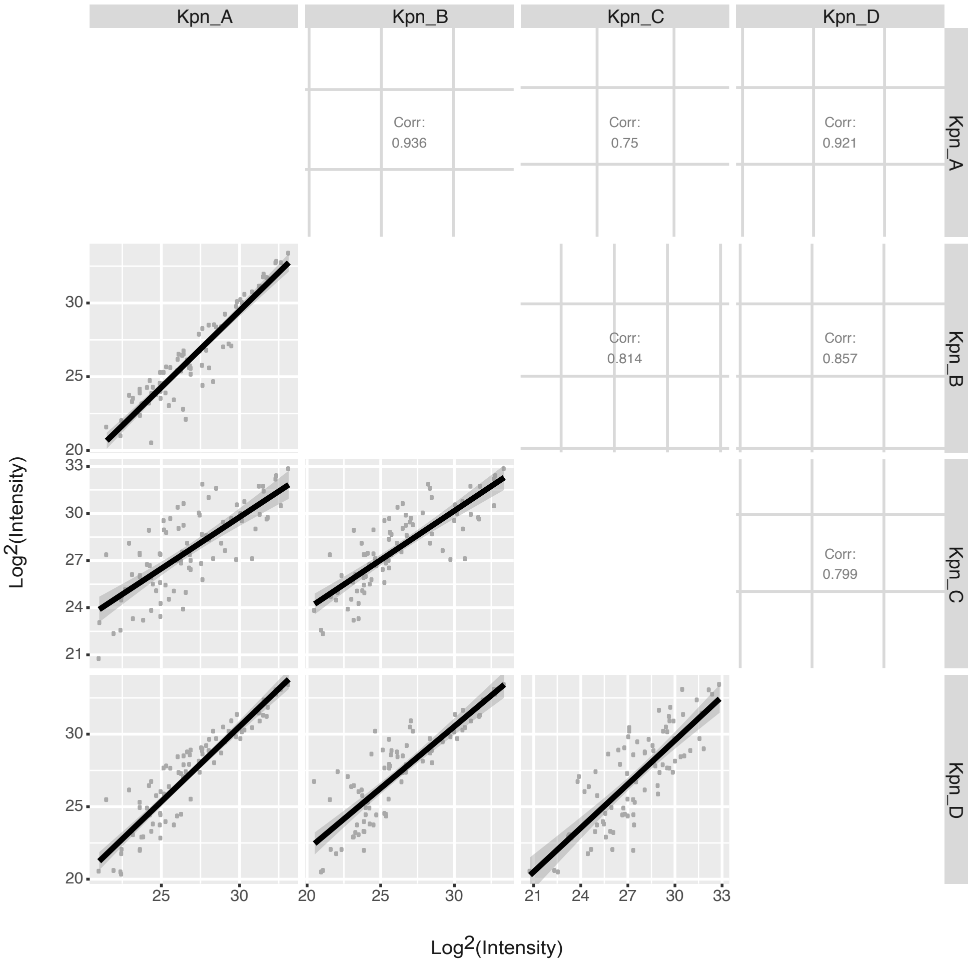


d

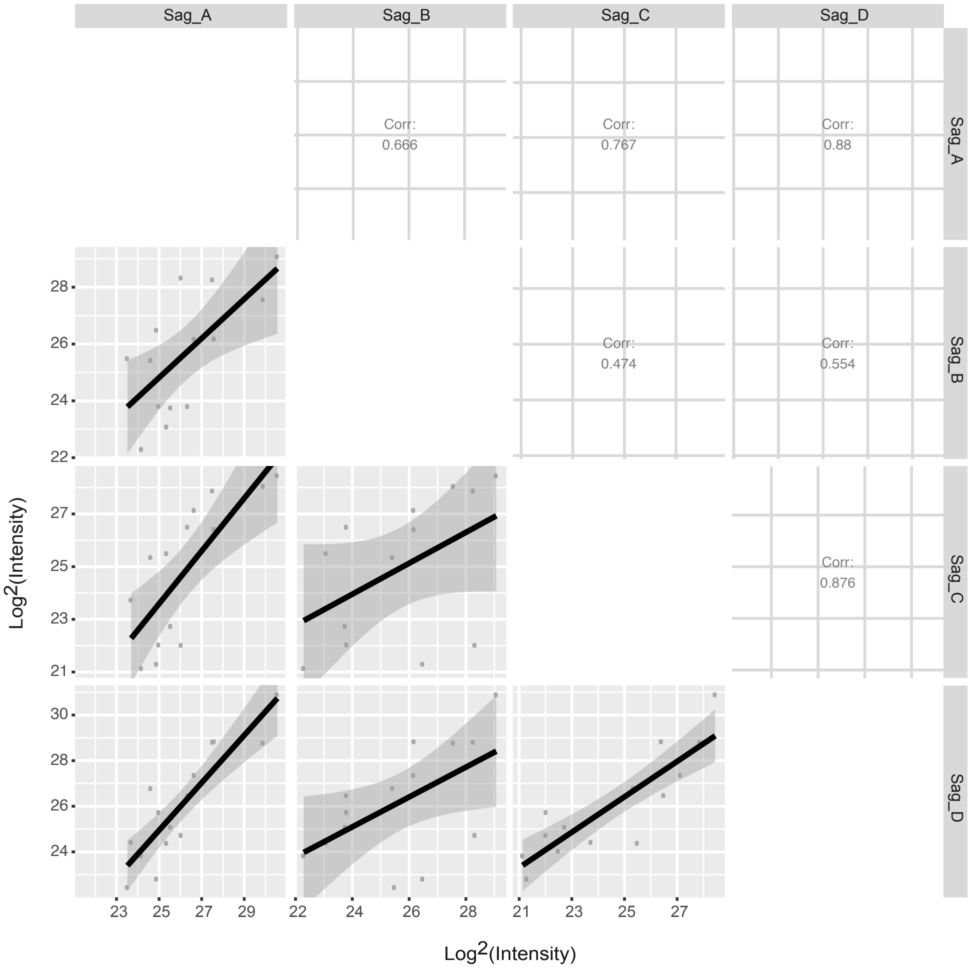

### Supplementary Table 1

| Organism | Gram | Taxon ID | Uniprot database Entries | Nbr Proteins | Nbr Protein Groups | Nbr Modified Peptide Groups | Proteome Coverage (%) | Total Genomic Length (Mb) | Predicted Protein Count |
|---|---|---|---|---|---|---|---|---|---|
| Citrobacter freundii (Cfr) | - | 546 | 5008 | 1806 | 1797 | 25705 | 36.1 | 5.27 | 4985 |
| Enterobacter cloacae (Ecl) | - | 550 | 4330 | 1134 | 1121 | 11407 | 26.2 | 4.93 | 4613 |
| Escherichia coli (Eco) | - | 83333 | 4314 | 1787 | 1772 | 24726 | 41.4 | 5.15 | 5009 |
| Klebsiella aerogenes (Kae) | - | 1028307 | 4909 | 1096 | 1085 | 10686 | 22.3 | 5.24 | 4915 |
| Klebsiella oxytoca (Kox) | - | 571 | 6408 | 1616 | 1583 | 17206 | 25.2 | 6.04 | 5608 |
| Klebsiella pneumoniae (Kpn) | - | 272620 | 5126 | 1914 | 1899 | 26888 | 37.3 | 5.59 | 5407 |
| Pseudomonas aeruginosa (Pae) | - | 208964 | 5564 | 2453 | 2438 | 29558 | 44.1 | 6.6 | 6103 |
| Proteus mirabilis (Pmi) | - | 584 | 3661 | 1307 | 1300 | 14913 | 35.7 | 3.95 | 3500 |
| Enterococcus faecalis (Efa) | + | 226185 | 3240 | 1128 | 1123 | 16986 | 34.8 | 2.99 | 2919 |
| Streptococcus agalactiae (Sag) | + | 208435 | 2105 | 867 | 863 | 13101 | 41.2 | 2.08 | 2015 |
| Staphylococcus aureus (Sau) | + | 93061 | 2889 | 1146 | 1142 | 15902 | 39.7 | 2.84 | 2820 |
| Staphylococcus epidermidis (Sep) | + | 176280 | 2461 | 1144 | 1140 | 13110 | 46.5 | 2.53 | 2373 |
| Staphylococcus haemolyticus (Sha) | + | 279808 | 2640 | 986 | 986 | 10987 | 37.3 | 2.5 | 2349 |
| Streptococcus mitis (Smi) | + | 28047 | 1983 | 810 | 810 | 11411 | 40.8 | 1.98 | 1838 |
| Staphylococcus saprophyticus (Ssa) | + | 342451 | 2404 | 1163 | 1161 | 14970 | 48.4 | 2.59 | 2401 |
| Total number of Peptide Groups | | | | | | 257556 | | | |
| Number of Non-Redundant Stripped Sequences | | | | | | 126859 | | | |
| Number of Non-Redundant Stripped Sequences after filtering | | | | | | 31096 | | | |

Supplementary Table 3

| Sample Name | Bacterial culture concentration (CFU/mL) | Inoculated volume (µL) | Final concentration in urine (CFU/mL) |
|---|---|---|---|
| For validation by targeted proteomics | | | |
| UP 1 | 4.63E+08 | 1 | 4.63E+04 |
| UP 2 | 4.63E+08 | 2 | 9.26E+04 |
| UP 10 | 4.63E+08 | 10 | 4.63E+05 |
| UP 20 | 4.63E+08 | 20 | 9.26E+05 |
| UP 100 | 4.63E+08 | 100 | 4.63E+06 |
| EF 1 | 8.77E+08 | 1 | 8.77E+04 |
| EF 2 | 8.77E+08 | 2 | 1.75E+05 |
| EF 10 | 8.77E+08 | 10 | 8.77E+05 |
| EF 20 | 8.77E+08 | 20 | 1.75E+06 |
| EF 100 | 8.77E+08 | 100 | 8.77E+06 |
| KP 1 | 6.30E+08 | 1 | 6.30E+04 |
| KP 2 | 6.30E+08 | 2 | 1.26E+05 |
| KP 10 | 6.30E+08 | 10 | 6.30E+05 |
| KP 20 | 6.30E+08 | 20 | 1.26E+06 |
| KP 100 | 6.30E+08 | 100 | 6.30E+06 |
| SA 2 | 1.28E+08 | 2 | 2.56E+04 |
| SA 4 | 1.28E+08 | 4 | 5.12E+04 |
| SA 20 | 1.28E+08 | 20 | 2.56E+05 |
| SA 40 | 1.28E+08 | 40 | 5.12E+05 |
| SA 200 | 1.28E+08 | 200 | 2.56E+06 |
| For comparison with MALDI-TOF | | | |
| Spike_Sag | 1.17E+08 | 100 | 1.17E+06 |
| Spike_Pae | 2.83E+08 | 200 | 5.66E+06 |
| Spike_Pmi | N/A | 20 | N/A |
| Spike_Ecl | 3.20E+08 | 20 | 6.40E+05 |
| Spike_Kox | 8.37E+08 | 20 | 1.67E+06 |
| Spike_Cfr | 8.40E+08 | 20 | 1.68E+06 |
| Spike_Kae | 9.30E+08 | 20 | 1.86E+06 |
| Spike_Sm | 3.97E+08 | 20 | 7.94E+05 |
| Spike_Ecc | 3.03E+08 | 20 | 6.06E+05 |
| Spike_Efa | 8.07E+08 | 20 | 1.61E+06 |
| Spike_Kpr | 6.07E+08 | 20 | 1.21E+06 |
| Spike_Sep | 1.17E+08 | 200 | 2.34E+06 |
| Spike_Ssa | 1.93E+08 | 20 | 3.86E+05 |
| Spike_Sau | 2.26E+08 | 100 | 2.26E+06 |
| Spike_Sha | 1.66E+08 | 20 | 3.32E+05 |

Supplementary Table 4

| Sample Name | MALDI-TOF | LC-MS |
|---|---|---|
| Patient urine specimens | | |
| Patient_038 | Escherichia coli | Eco |
| Patient_C01 | Absence of uropathogens | Blk |
| Patient_C04 | Staphylococcus saprophyticus | Ssa |
| Patient_C06 | Absence of uropathogens | Blk |
| Patient_C07 | Streptococcus epidermidis | Sep |
| Patient_C08 | Absence of uropathogens | Blk |
| Patient_C10 | Absence of uropathogens | Blk |
| Patient_C11 | Escherichia coli | Eco |
| Patient_C12 | Absence of uropathogens | Blk |
| Patient_C14 | Streptococcus probable | Blk |
| Patient_D02 | Enterococcus problable | Blk |
| Patient_D03 | Enterococcus problable | Blk |
| Patient_D05 | Escherichia coli | Eco |
| Patient_D10 | Streptococcus agalactiae | Blk |
| Patient_D14 | Escherichia coli | Eco |
| Patient_D21 | Escherichia coli | Eco |
| Patient_D22 | Staphylococcus probable | Blk |
| Patient_D26 | Staphylococcus probable | Blk |
| Patient_D31 | Escherichia coli | Eco |
| Patient_D32 | Staphylococcus probable | Blk |
| Patient_D40 | Klebsiella pneumoniae | Kpn |
| Patient_D47 | Escherichia coli | Kae |
| Patient_E01 | Absence of uropathogens | Blk |
| Patient_E05 | Absence of uropathogens | Blk |
| Patient_E08 | Absence of uropathogens | Blk |
| Patient_E09 | Escherichia coli | Eco |
| Patient_E30 | Escherichia coli | Eco |
| Patient_E34 | Enterococcus problable | Blk |
| Patient_E39 | Enterococcus faecalis | Efa |
| Patient_E42 | Escherichia coli | Eco |
| Patient_E48 | Escherichia coli | Eco |
| Bacterial inoculation in urine | | |
| Spike_Cfr | Citrobacter freundii | Cfr |
| Spike_Ecl | Enterococcus cloacae complex | Ecl |
| Spike_Eco | Escherichia coli | Eco |
| Spike_Efa | Enterococcus faecalis | Efa |
| Spike_Kae | Enterococcus cloacae complex | Kae |
| Spike_Kox | Klebsiella oxytoca | Kox |
| Spike_Kpn | Klebsiella pneumoniae | Kpn |
| Spike_Pae | Pseudomonas aeruginosa | Pae |
| Spike_Pmi | Proteus mirabilis | Pmi |
| Spike_Sag | Streptococcus agalactiae | Sag |
| Spike_Sau | Staphylococcus aureus | Sau |
| Spike_Sep | Streptococcus epidermidis | Sep |
| Spike_Sha | Staphylococcus haemolyticus | Sha |
| Spike_Smi | Candida albicans | Smi |
| Spike_Ssa | Staphylococcus saprophyticus | Ssa |
